# Development of a 1:1-binding biparatopic anti-TNFR2 antagonist by epitope selection

**DOI:** 10.1101/2022.12.15.520217

**Authors:** Hiroki Akiba, Junso Fujita, Tomoko Ise, Kentaro Nishiyama, Tomoko Miyata, Takayuki Kato, Keiichi Namba, Hiroaki Ohno, Haruhiko Kamada, Satoshi Nagata, Kouhei Tsumoto

**Affiliations:** Graduate School of Pharmaceutical Sciences, Kyoto University; Sakyo-ku, Kyoto, 606-8501, Japan; Collaborative Research Center for Health and Medicine, National Institutes of Biomedical Innovation, Health and Nutrition; Ibaraki City, Osaka, 562-0011, Japan; Graduate School of Frontier Biosciences, Osaka University; Suita City, Osaka, 565-0871, Japan; Graduate School of Pharmaceutical Sciences, Osaka University; Suita City, Osaka, 565-0871, Japan; Center for Drug Design Research, National Institutes of Biomedical Innovation, Health and Nutrition; Ibaraki City, Osaka, 562-0011, Japan; JEOL YOKOGUSHI Research Alliance Laboratories, Osaka University; Osaka, 565-0871, Japan; Institute of Protein Research, Osaka University; Suita City, Osaka, 565-0871, Japan; RIKEN SPring-8 Center; Osaka 565-0871, Japan; School of Engineering, The University of Tokyo; Bunkyo-ku, Tokyo, 113-8656, Japan; Institute of Medical Sciences, The University of Tokyo; Minato-ku, Tokyo, 108-8639, Japan

## Abstract

Conventional bivalent antibodies against cell surface receptors often initiate unwanted signal transduction by crosslinking two antigen molecules. Biparatopic antibodies (BpAbs) bind to two different epitopes on the same antigen, thus altering crosslinking ability. In this study, we developed BpAbs against tumor necrosis factor receptor 2 (TNFR2), which is an attractive immune checkpoint target. Using different pairs of variable regions specific to topographically distinct TNFR2 epitopes, we successfully regulated the size of BpAb-TNFR2 immunocomplexes to result in controlled agonistic activities. One particular antagonist BpAb bound TNFR2 in 1:1 ratio without unwanted signal transduction, with its structural basis revealed by cryo-electron microscopy. This antagonist suppressed the proliferation of regulatory T cells expressing TNFR2. Therefore, the BpAb format would be useful in designing specific and distinct antibody functions.

## Introduction

Tumor necrosis factor receptor 2 (TNFR2) is a member of the tumor necrosis factor receptor superfamily (TNFRSF) that cluster on the cellular membrane by interacting with their specific ligands (*1-3*). TNFR2 aggregates upon binding to trivalent tumor necrosis factor alpha (TNFα) (Fig. 1A,B) (*1-4*). Aggregated TNFR2 increases the density of receptor-associated intracellular proteins, triggering the activation of downstream canonical and noncanonical NF-κB pathways (*1,5,6*). TNFR2 is primarily expressed in a subset of T cells, including regulatory T cells (T_regs_) (*7-10*), and TNFR2-mediated intracellular signaling expands T_regs_, inducing cancer cell proliferation. Therefore, anti-TNFR2 antagonists are promising as a new type of immune checkpoint inhibitor to enhance the immune response in patients with cancer (*9-12*). In contrast, anti-TNFR2 signal-inducing antibodies (agonists) are pursued as therapeutic strategies to expand T_regs_ in autoimmunity, organ transplant rejection, and graft-versus-host disease. (*9,10,13-15*).

**Fig. 1.**
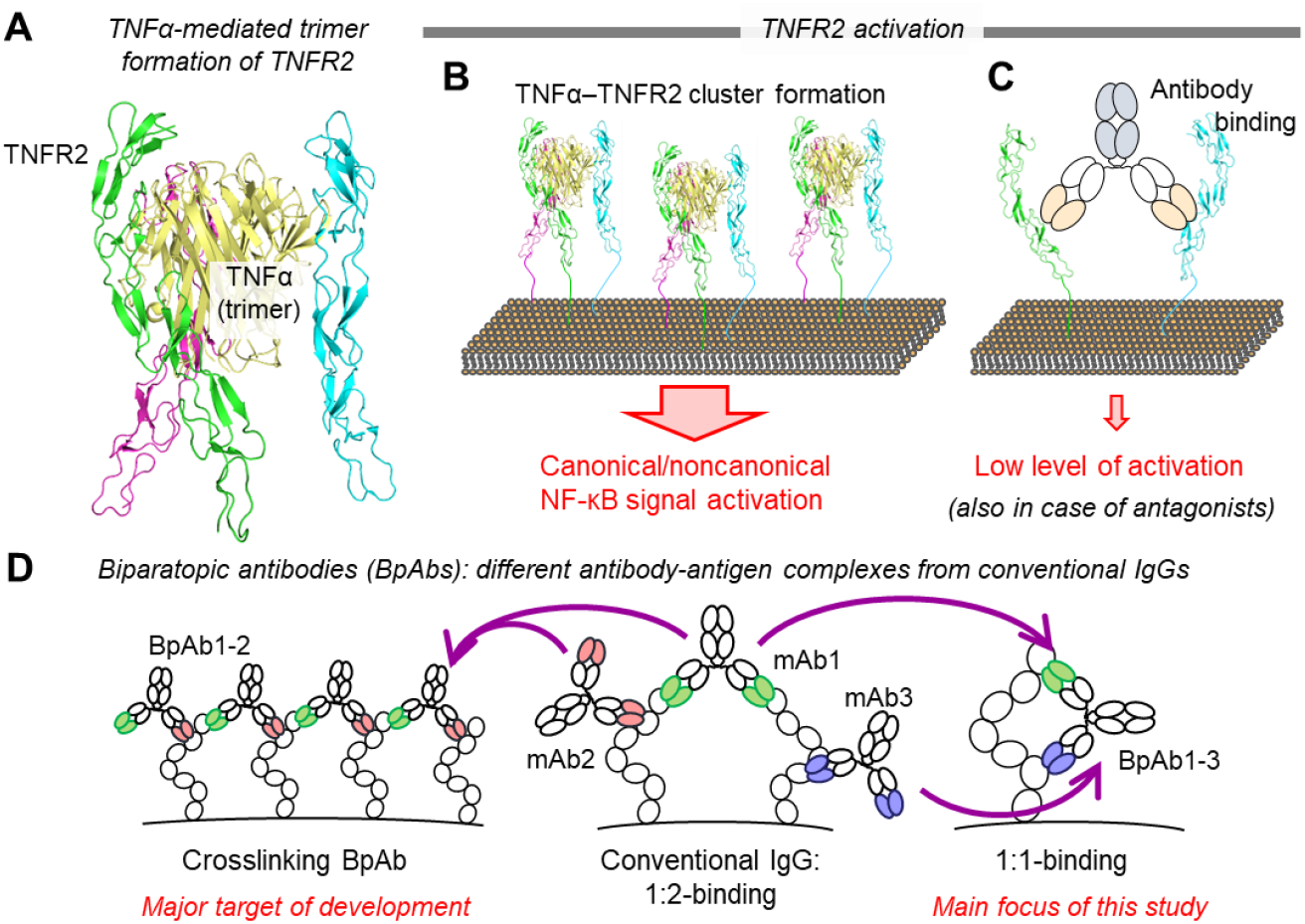
Activation of TNFR2 and biparatopic antibodies (BpAbs). (**A**) TNFR2 trimer is formed by interaction with TNFα (PDB ID: 3ALQ). (**B**) Trimer formation induces cluster formation of TNFR2 leading to activation of NF-κB signaling pathways. (**C**) Conventional antibodies, including antagonists, bind two TNFR2 molecules and may activate low levels of signaling. (**D**) Conventional IgG antibody binds the antigen in 1:2 manner, but BpAbs including crosslinking type of BpAbs (BpAb1-2) and 1:1-binding BpAbs (BpAb1-3) involve a different topology of antibody-antigen complexes compared to conventional IgGs.

Bivalency of conventional antibody (cIgG) could produce dimer formation-related unwanted cellular responses (*16*). Similar to the natural TNFα ligand (Fig. 1B), anti-TNFR2 cIgGs may induce TNFR2 clustering by bivalent binding (Fig. 1C). Thus, the agonistic activity of antagonist antibodies needs to be minimized to widen their clinical application potential. A conventional antagonistic antibody requires a specialized binding mode related to specific antigen-binding sites (epitopes) to prevent functional clustering (*11*). However, this binding mode depends on empirical functional screening, thus hindering the design of an effective antagonist. In this study, we developed biparatopic antibodies (BpAbs) against TNFR2 to enable a more effective design of antagonists. BpAb is an engineered bispecific antibody bearing two different antibody variable fragments (Fv) that bind to distinct epitopes on the same antigen (Fig. 1D). By characterizing multiple bivalent IgG-like BpAbs targeting different topographical TNFR2 epitopes, we demonstrated that variable levels of agonistic activity could be achieved by bivalent BpAbs, including BpAbs with no agonistic activity. The agonistic potency clearly depended on the ability of cIgGs and BpAbs to form clustered immunocomplexes with the antigen. We demonstrated the binding of one potential antagonist, Bp109-92, to TNFR2 in 1:1 manner by physicochemical characterization and cryo-electron microscopy (cryo-EM) (Fig. 1D, right). We showed that the functional activities of the produced BpAbs are predictable based on the relative positions of the two epitopes. Thus, our findings lead to a fine-tuned design of both agonists and antagonists based on BpAbs.

## Results

### Comprehensive production and biological activities of biparatopic antibodies

We identified five monoclonal antibodies (mAbs) binding different epitopes of TNFR2 (TR45, TR92, TR94, TR96, and TR109), which were confirmed by competitive ELISA (Fig. 2A). The binding sites revealed by multiple mutation studies are mapped in Fig. 2B and 2C. To identify epitope sites on the surface of TNFR2, we designed a series of TNFR2 mutants by peptide grafting (Fig. S1A and Table S1). The five anti-TNFR antibodies used in this study (TR45, TR92, TR94, TR96, and TR109) were selected from an epitope-normalized antibody panel, which is a technology to obtain a panel of a minimum number of antibodies whose epitope regions are comprehensively and evenly distributed over the entire accessible surface of a target antigen (WO2018/092907). First, mutants were generated to reduce binding to human TNFR2. For each mutant, the respective region of human TNFR2 was substituted with that of mouse TNFR2 (human-to-mouse mutants). The extracellular region of TNFR2 contained four cysteine-rich domains (CRDs; CRD1, 2, 3 and 4; Fig. S1B). As a first set of mutants, CRD1, 2 and 3 were split by half, and added CRD4 and C- and N-terminal region substituents to produce nine mutants (‘Mut1’ line of Fig. S1A; Fig. S1C). We produced nine additional mutants (‘Mut2’ line in Fig. S1A; Fig. S1D) for detailed analysis. These human-to-mouse mutants were named mC1A. Epitope sites were identified by reduced binding to cells transiently transfected with wild-type human TNFR2 or human-to-mouse mutants, followed by analysis using a flow cytometer. Regions with total or partial loss of binding compared to the wild-type were identified (Fig. S2,S3). As a complementary analysis, mouse-to-human mutants were produced by replacing the peptides in mouse TNFR2 with a human ortholog (referred to as hC1A). Binding was only observed when the epitope region was present (Fig. S4,S5), and TR109 required both hC1A2 and hC2A2 regions for binding. This result was consistent with the analysis of reduced binding and supported the epitope identification analysis. All five mAbs bound to different parts of the human TNFR2. These regions were mapped onto structures (Fig. 1B). The epitopes of TR92 and TR109 overlap with the TNFα-binding region. The epitope of TR45 was non-overlapping but on the same side as the TNFα-binding region, whereas those of TR94 and TR96 were located on the opposite side.

**Fig. 2.**
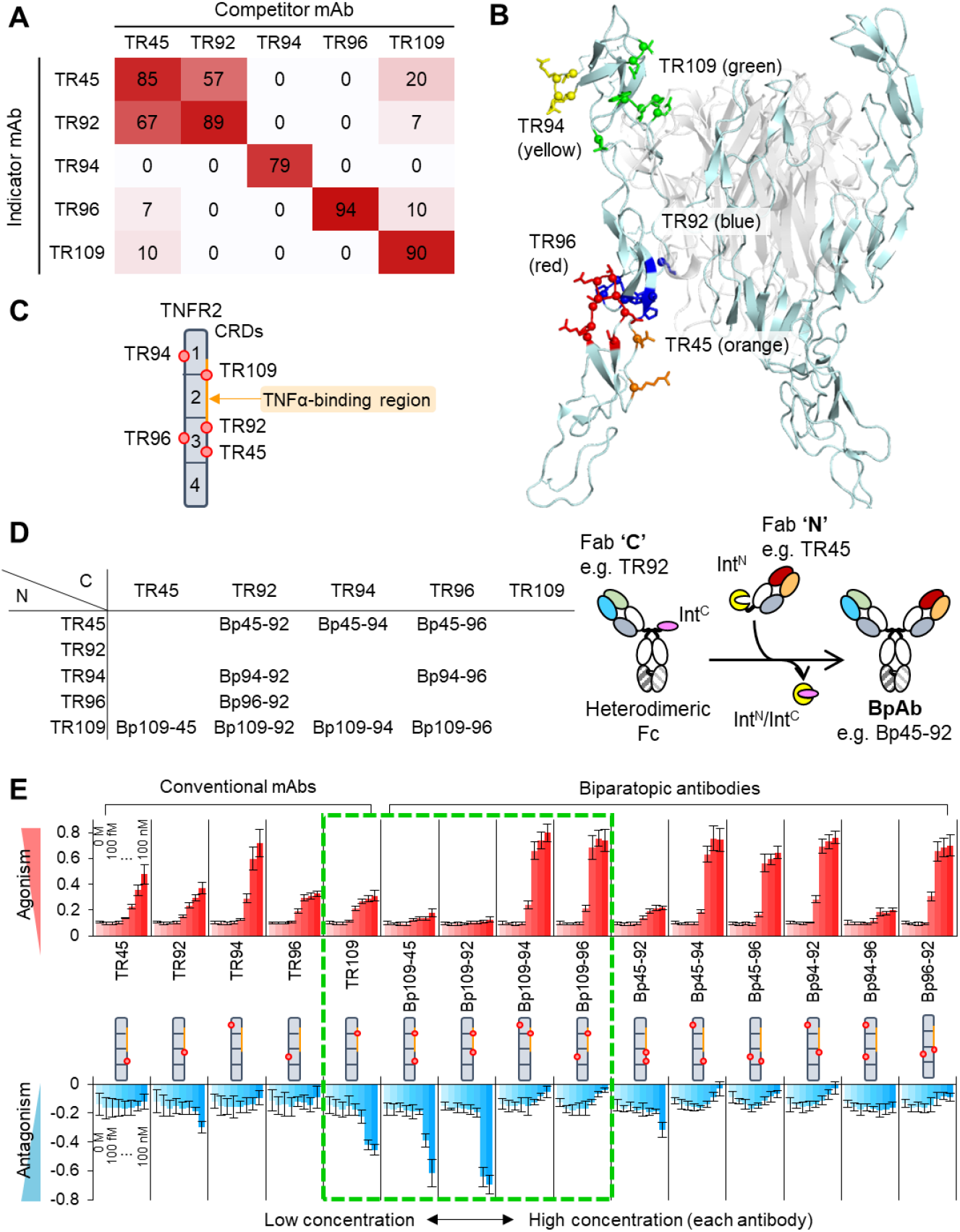
Monoclonal antibodies and biparatopic antibodies against TNFR2. (**A**) Topographical epitope mapping of the anti-TNFR2 mAbs by mutual competition analysis. Strengths of competition are shown as percentages in each box. (**B**) Epitopes of the selected monoclonal antibodies identified using the mutants. Cyan, TNFR2; white, TNFα (PDB ID: 3ALQ). (**C**) Schematic presentation of the epitopes (red circles). Cysteine-rich domains (CRDs) are separated by lines. (**D**) Produced BpAbs. Fab ‘**N**’ (row) and ‘**C**’ (column) were fused with Int^N^ or Int^C^-Fc, respectively, and were subjected to intein-mediated protein trans-splicing to produce a BpAb (*31*). (**E**) Agonistic (upper) and antagonistic (lower) activity of conventional and biparatopic antibodies, monitored using a reporter cell line. Values are shown with standard deviation of five independent experiments standardized by the reporter activity of 0 and 50 ng/mL TNFα.

Bivalent IgG-like BpAbs of all ten possible combinations of Fvs from the five mAbs were produced (Fig. 2D), and confirmed to bind to the same epitopes as the original cIgGs (Fig. S4,S5).

Next, we used reporter cells to detect NFκB-dependent DNA transcription in the absence or presence of TNFα to evaluate the agonistic and antagonistic activities of cIgGs and BpAbs (Fig. 2E). The BpAb panel included the majority of six strong agonists with high maximum agonistic activity at low concentrations compared with those of cIgGs. This is consistent with reported BpAbs bridging multiple antigen molecules and utilized as agonists for cell-surface proteins, including TNFRSF (Fig. 1D, left) (*17-20*). It should be noted that all five cIgGs were moderately agonistic, as predicted based on their bivalency. In contrast, BpAbs contained two antagonists with significantly reduced agonistic activity (Bp109-45 and Bp109-92). Bp109-92 displayed a strong antagonistic effect against TNFα-dependent activity without any apparent agonistic activity. Thus, we successfully produced an artificial antagonistic BpAb that has previously never been generated naturally. Altogether, these results suggest that BpAbs can be designed with desired functions using selected pairs of Fvs that recognize two non-overlapping epitopes.

### Binding analysis and the size of immunocomplexes

We analyzed the concentration-dependent binding of cIgGs and BpAbs to reporter cells using flow cytometry to uncover the origin of their agonistic activities. All cIgGs and BpAbs, except for TR94, showed a typical sigmoidal curve, with an IC_50_ of approximately 30 ng/mL (corresponding to 0.2 nM) (Fig. S6). The affinity of cIgGs and BpAbs for TNFR2 was also measured by surface plasmon resonance. Most BpAbs bound more strongly than cIgGs (*K*_D_ = 0.92 nM versus 7.7 nM on average; Fig. S7 and Table S2). In either experiment, the strength of the interaction did not correlate with maximum agonistic activity, however the curved shape of cellular binding coincided with the agonistic activity (Fig. S8). The binding-activity relationship indicated that the higher-order structure of the immunocomplex was key for determining functional activity.

Size-exclusion chromatography with multi-angle light scattering (SEC-MALS) was conducted to analyze the size of the antigen-antibody complexes. Chromatograms of TR109 and the three BpAbs using the variable regions of TR92, TR96, and TR109 are shown in Fig. 3A. TNFR2 had a molar mass of 37 kDa, and Bp96-92 alone had a molar mass of 150 kDa (other cIgGs and BpAbs were of the same size; Fig. S9). TR109 formed a homogeneous 1:2 complex of antibody:TNFR2 of 220 kDa in the presence of excess TNFR2. Bp96-92 and Bp109-96 formed larger heterogeneous complexes, with a broad distribution of molar masses ranging from 200 to 500 kDa. These two BpAbs were capable of forming large bridging immunocomplexes. In contrast, Bp109-92 formed a homogeneous 1:1 complex of 190 kDa in the presence of excess TNFR2; thus, 1:1 complex formation was more advantageous than 1:2 complex formation. The size of the complex was analyzed using mass photometry at the single-particle level (*21*) and similar results were obtained. For Bp109-96 and Bp96-92, equimolar BpAb-antigen complexes up to an antibody:TNFR2 ratio of 4:4 were observed (Fig. 3B,C). In contrast, the antibody:TNFR2 1:1 complex was dominant in Bp109-92 (Fig. 3D). The observation of equimolar complexes indicated that the two variable regions of BpAb were both bound to TNFR2, and the observation by SEC-MALS was the average of mixed immunocomplexes at elution, reflecting their size distribution quantitatively.

**Fig. 3.**
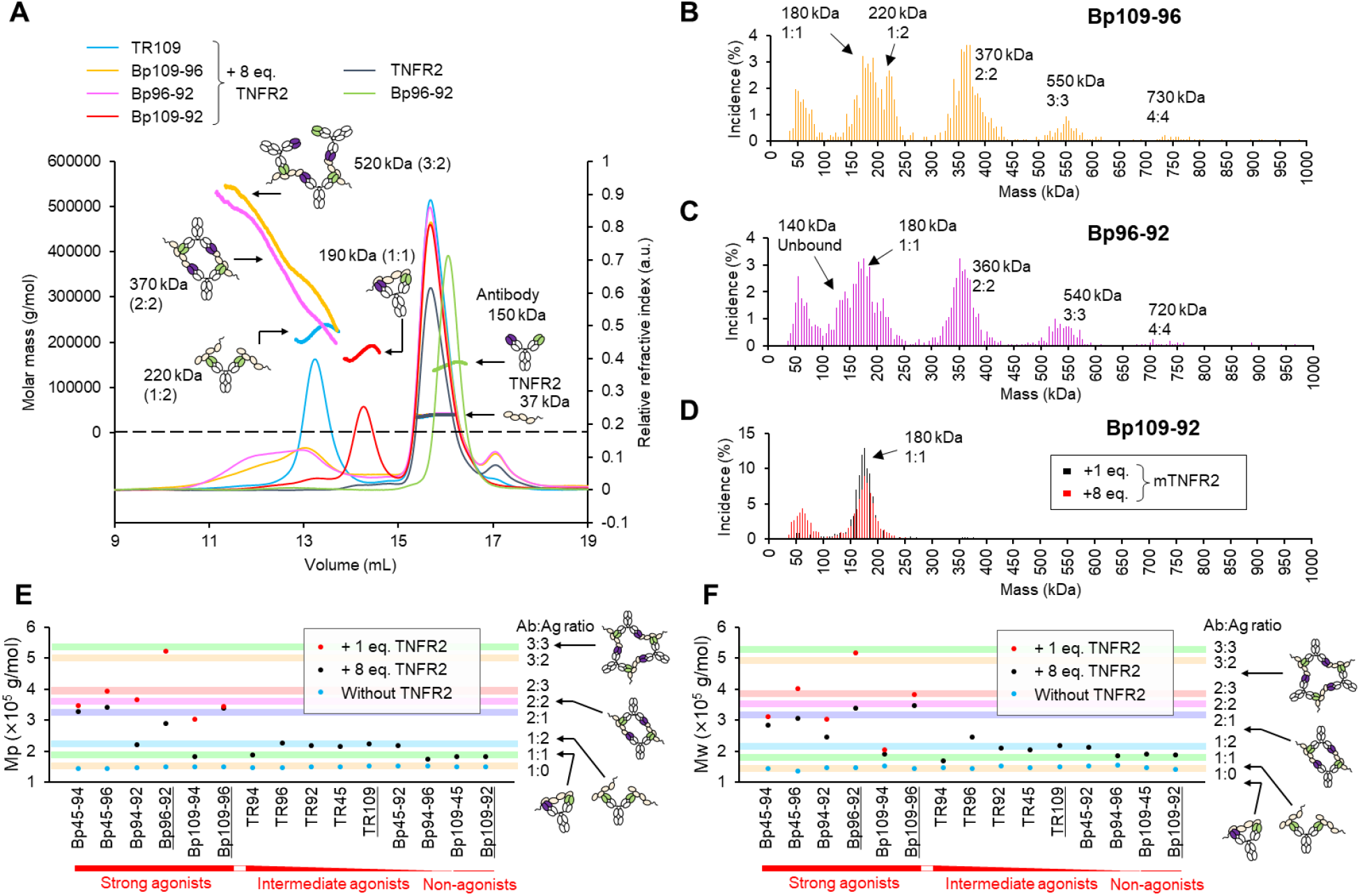
Size of the immunocomplexes formed by conventional and biparatopic antibodies analyzed by SEC-MALS (A,E,F) and mass photometry (B–D). (**A**) Selected chromatograms of SEC-MALS. Relative refractive index and molar mass are shown with thin and bold lines, respectively. (**B–D**) Relative frequency of the observed particles with the respective mass in mass photometry. Bp109-96 (**B**), Bp96-92 (**C**) or Bp109-92 (**D**) was mixed with 1 eq. or 8 eq. (only for Bp109-92) of TNFR2. (**E**,**F**) Peak molecular weight (Mp) (**E**) or weight-averaged molecular weight (Mw) (**F**) of the total fractions of immunocomplexes (eluted earlier than 14.8 mL). Chromatograms of underlined antibodies are shown in (**A**). Colored bands represent the estimated size of the immunocomplex with the expected structures in the right column.

Other cIgGs and BpAbs were also analyzed using SEC-MALS (Fig. 3E,F, and S9). All six strong agonist BpAbs formed large immunocomplexes, whereas the immunocomplexes of the non-agonist BpAbs were smaller than those of cIgGs. As an exception, poor binding ability of TR94 (Fig. S7) resulted in a smaller complex size. A similarly smaller immunocomplex was observed for Bp109-94 in the presence of excess TNFR2, but a large immunocomplex was observed when equimolar amounts of TNFR2 were mixed (Fig. S9). The larger immunocomplex observed for the equimolar mixture was shared among agonistic bridging BpAbs, possibly due to the disadvantage of 1:2 complex formation (Fig. S10). In summary, cIgGs and BpAbs were categorized into three classes: (1) large immunocomplexes formed by strong agonist BpAbs, (2) 1:2 complex of antibody:TNFR2 by moderate agonists, and (3) 1:1 immunocomplexes without agonistic activity including antagonistic BpAbs. A simple relationship was observed over a broad concentration range.

Bp45-94 (agonist) and Bp109-92 (antagonist) in complex with TNFR2 were analyzed using negative-stain EM. In both cases, TNFR2 was not visible owing to its small size and low resolution. Bp45-94 showed a variety of structures up to a complex comprising three BpAb molecules; however, most particles were monomers or dimers (Fig. S11A). Monomers were presumably unbound or bound by one or two TNFR2 molecules but were not distinguishable. The dominant larger complex was a dimer, consistent with the observations by SEC-MALS. On the other hand, mostly monomers were observed for Bp109-92, even in the presence of TNFR2 (Fig. S11B). When the Bp109-92 F(ab′)_2_–TNFR2 complex was analyzed by enzymatic removal of Fc, a parallel arrangement of two antigen-binding fragments (Fabs) was observed (Fig. S12A). This is in contrast to the random orientation of Bp109-92 F(ab′)_2_ in the absence of TNFR2 (Fig. S12B).

### Precise characterization of the antagonist, Bp109-92

After several rounds of optimization, the three-dimensional structure of the Bp109-92– TNFR2 1:1 complex was determined at a resolution of 3.63 Å by cryo-EM single-particle image analysis using a TNFR2-maltose binding protein fusion as part of the complex (Fig. S13–14, Table S3). In the complex structure (Fig. 4A), Fabs from both TR92 and TR109 (92-Fab and 109-Fab) were in a parallel arrangement, as observed by negative-stain EM (Fig. S12), and biparatopic binding involving the simultaneous binding of both variable regions was observed. The epitope locations of the two mAbs determined by mutagenesis matched the Fab–TNFR2 interfaces (Fig. 4B,C), which was shared by the TNFR2–TNFα interface in the reported crystal structure (PDB ID:3ALQ) (Fig. 4D) (*4*). The density corresponding to Fc was not clearly observed. As for the antigen, TNFR2 possesses 4 cysteine rich domains (CRDs) as structure units in the extracellular region. The atomic model of TNFR2 was only partially built, ranging from cysteine-rich domain 1 (CRD1) to the middle of CRD3, corresponding to the C3A region in Fig. S1, although the density corresponding to CRD3 and CRD4 was weak (Fig. S15). CRD3 and CRD4 were oriented differently from the crystal structure of TNFR2 in complex with TNFα (Fig. 4D and Fig. S17A). The overall structures of 109-Fab and 92-Fab were similar (r.m.s.d. = 1.20 Å among 397 C_α_ atoms), but the CDR regions were significantly changed to interact with different regions of TNFR2 (Fig. S16). In the TNFR2–TNFα complex, Gln109 of TNFR2 interacts with TNFα, and the loop of the C3A2 region (orange in Fig. 4D) is pulled toward TNFα. In contrast, this loop partly protruded into a unique cleft between heavy chain complementarity-determining regions 1 (CDR-H1) and CDR-H3 of 92-Fab in the Bp109-92 complex (Fig. 4C) and pulled in the opposite direction. Leu118 at the C-terminus of C3A was shifted by 13 Å when the two TNFR2 structures were aligned by CRD1 and CRD2 (Fig. 4D). Flexibility of the CRDs of TNFR2 was indicated.

**Fig. 4.**
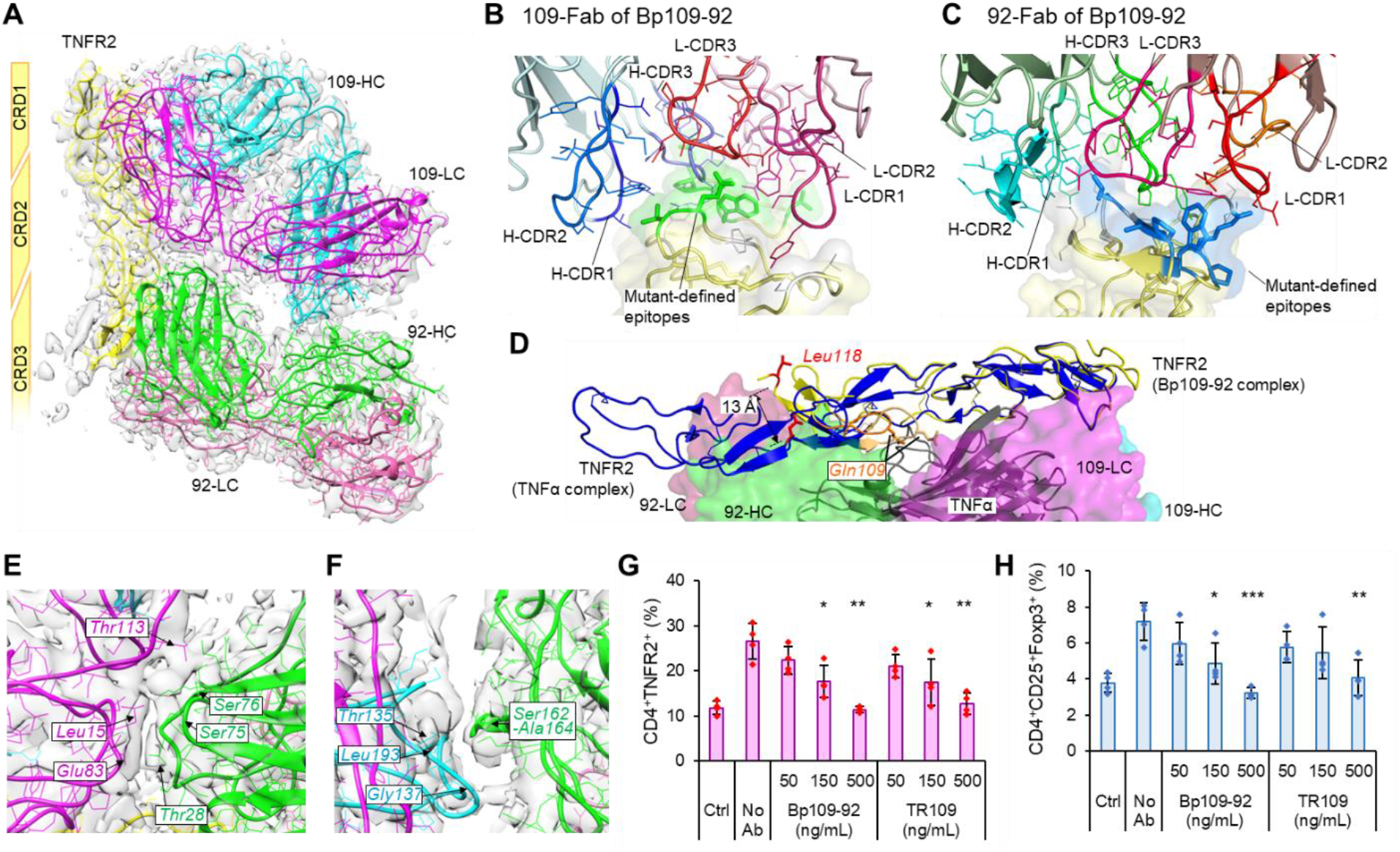
(A–F) Cryo-EM structure of Bp109-92 in complex with TNFR2. (**A**) Overall view of the cryo-EM structure. 109-HC/LC and 92-HC/LC are the heavy/light chains from TR109 and TR92, respectively. (**B**,**C**) Enlarged view of the epitope site of 109-Fab (**B**) and 92-Fab (**C**) observed for the complex with BpAb. Each complementarity-determining region (CDR) loop is colored separately. Mutant-defined epitope amino acids are colored green (**B**) or blue (**C**), while interacting amino acids not clearly defined as the epitope residue are colored white. (**D**) Structural difference of TNFR2 observed in complex with Bp109-92 (yellow) and with TNFα (blue) (PDB ID: 3ALQ). 109-Fab and 92-Fab are shown with the van der Waals surface in the same color as (**A**), while TNFα is shown with cartoon in grey. Leu118 and C3A2 region of the two TNFR2 structures are colored red and orange, respectively. (**E**,**F**) Fab-Fab contacts inside Bp109-92 in complex with TNFR2. (**E**) Interface between 109-LC and 92-HC. (**F**) Interface between 109-HC and 92-HC. **(G**,**H) Suppression of TNFR2 activity of human T cells by the antagonists**. Population of CD4^+^TNFR2^+^ cells (**G**) or CD4^+^CD25^+^Foxp3^+^ cells (**H**) among CD3^+^ cells in PBMC without treatment (Ctrl), in the presence of 100 ng/mL TNFα (No Ab), or in the presence of TNFα and antibodies. The values obtained in each experiment are shown as dots, and the average of four experiments are shown with bars. Error bars represent the standard deviations. *Student’s t-test in comparison with ‘No Ab’, *, *P* < 0.05; **, *P* < 0.005; ***, *P* < 0.0005.

For Bp109-92 bound to TNFR2, contact between 109-Fab and 92-Fab was observed (Fig. 4E,F). This contact surface was formed by the heavy chain variable domain (VH) of 92-Fab entering the concave surface at the elbow region of 109-Fab and interacting with the linker between the variable and constant regions of the light chain. Although the resolution was not high enough to discuss the respective atomic-level contacts, this may contribute energetically to the stabilization of the immunocomplex.

Finally, we used peripheral blood mononuclear cells (PBMC) to demonstrate the applicability of Bp109-92 in the inhibition of TNFα-dependent proliferation of T cells (Fig. S17 for the gating strategy). TNFα stimulation increased the population of TNFR2^+^ cells among CD3^+^ T cells (Fig. S18). This population coincided with CD4^+^CD25^+^CD127^low^ cells, which contain Foxp3^+^ regulatory T cells (Fig. S19). Both TR109 or Bp109-92 effectively reduced TNFR2^+^ cells, thus acting as antagonists of TNFα-dependent proliferation (Fig. 4G). The population of TNFR2^+^ cells increased from 12% to 27% after TNFα treatment; however, 500 ng/mL Bp109-92 or TR109 suppressed the population to 11% and 13%, respectively. A similar reduction was observed for Foxp3^+^ cells, which were also increased by TNFα (Fig. 4H). The population of Foxp3^+^ cells increased from 3.8% to 7.2% with TNFα treatment, which was reduced to 3.2% and 4.1% in the presence of 500 ng/mL Bp109-92 or TR109, respectively. Thus, both Bp109-92 and TR109 were effective antagonists, and the antagonistic activity of Bp109-92 was greater than that of TR109.

## Discussion

Recent advances in antibody engineering have paved the way to regulate the stoichiometry and size of complexes between antigen and antibody molecules (*22*). As one of the most promising techniques, artificial BpAbs have overcome the limitations of cIgGs by releasing the restriction of bivalency (*17-20*). Manipulating the size of the complex enables BpAbs against cell-surface receptors to control intracellular signaling on a mechanistic basis. Nevertheless, rational selection of appropriate pairs of Fvs for designing desired functional BpAbs has not yet been established. In the present study, we demonstrated that the signal transduction activities of anti-TNFR2 BpAbs were precisely controlled by the epitope regions used in bivalent BpAbs. Such clear switching of agonism and antagonism of BpAbs was not achieved in the cIgG format. Furthermore, the use of a bivalent IgG-like bispecific format has enabled a variety of signaling activities. Notably, 1:1 complex formation by Bp109-92 resulted in almost complete elimination of the agonistic activity and maximized the antagonistic function. This type of 1:1 binding complex is similarly prepared using single Fab-based formats (*23*). However, a single Fab lacks binding avidity and its binding activity is reduced. In contrast, the affinity of Bp109-92 for TNFR2 was 25- and 35-fold higher than parental cIgGs (Table S2), and similarly strong binding as the parental cIgGs with avidity was achieved (Fig. S6). Increased apparent binding affinity by designing BpAbs is common (*19*), including the case of estimated 1:1 binding with trimeric viral antigens (*24,25*). Although 1:1 binding may not be required for the biological activities of these antibodies, our results provide the first evidence that the exclusive formation of a specific 1:1 complex is required for biological significance. Thus, we showed that limiting immunocomplexes to a 1:1 ratio with cell-surface proteins can be an effective therapeutic approach.

Agonistic activity dependent on complex size can be controlled simply by the relative position of the epitopes of the two variable regions used for BpAb. Schematically, the CRDs of TNFR2 constituted a cylinder-like structure, and the epitopes of the five mAbs were mapped onto the cylindrical model viewed from the top (Fig. 5A). The epitopes of the five mAbs are grouped into group I consisting of the epitopes of TR109, TR92, and TR45 on the same side as the TNFα-binding region of TNFR2, and group II consisting of the other two epitopes. In the top view of TNFR2, epitopes in the same group were in proximity, and the two Fabs formed acute angles with each other (Fig. 5B, left), as observed in the cryo-EM structure of Bp109-92 (Fig. 4A). BpAbs designed by selecting two epitopes in the same group would be advantageous for forming a 1:1 complex of antibody and antigen unless competition occurs. On the other hand, when epitopes from two different groups are not proximal enough (Fig. 5B, right), 1:1 complex formation is impossible and large n:n complexes (n=2,3,4…) would dominate. Once the immunocomplex is formed, 1:1 complexes do not induce association-dependent signal transduction (non-agonist), 1:2 complexes formed by cIgGs work as weak agonists, and large n:n complex formation induces strong signal transduction (strong agonist). As seen in the cryo-EM structure of the Bp109-92– TNFR2 complex, the CRDs of TNFR2 may have some flexibility; thus, both 1:1 and large n:n complex formation would be possible without steric or torsional hindrance. The epitope dependency of the agonistic activity of BpAbs against TNFR2 in our study would guide the development of both effective agonists and antagonists against other members of TNFRSF proteins that share similar activation mechanisms. Following tetravalent BpAbs known to agonize OX40 effectively (*17*), the design contributes to the production of molecular therapeutics for immunology and oncology (*3,16,26,27*). The design, however, may depend on the mechanism of TNFRSF activation. For example, receptor tyrosine kinases are activated by phosphorylation, which is activated by the formation of a specific dimerized structure. In these cases, specific BpAbs or alternative scaffolds are designed to lock them into inactive structures (*28-30*), and more precise design may be required.

**Fig. 5.**
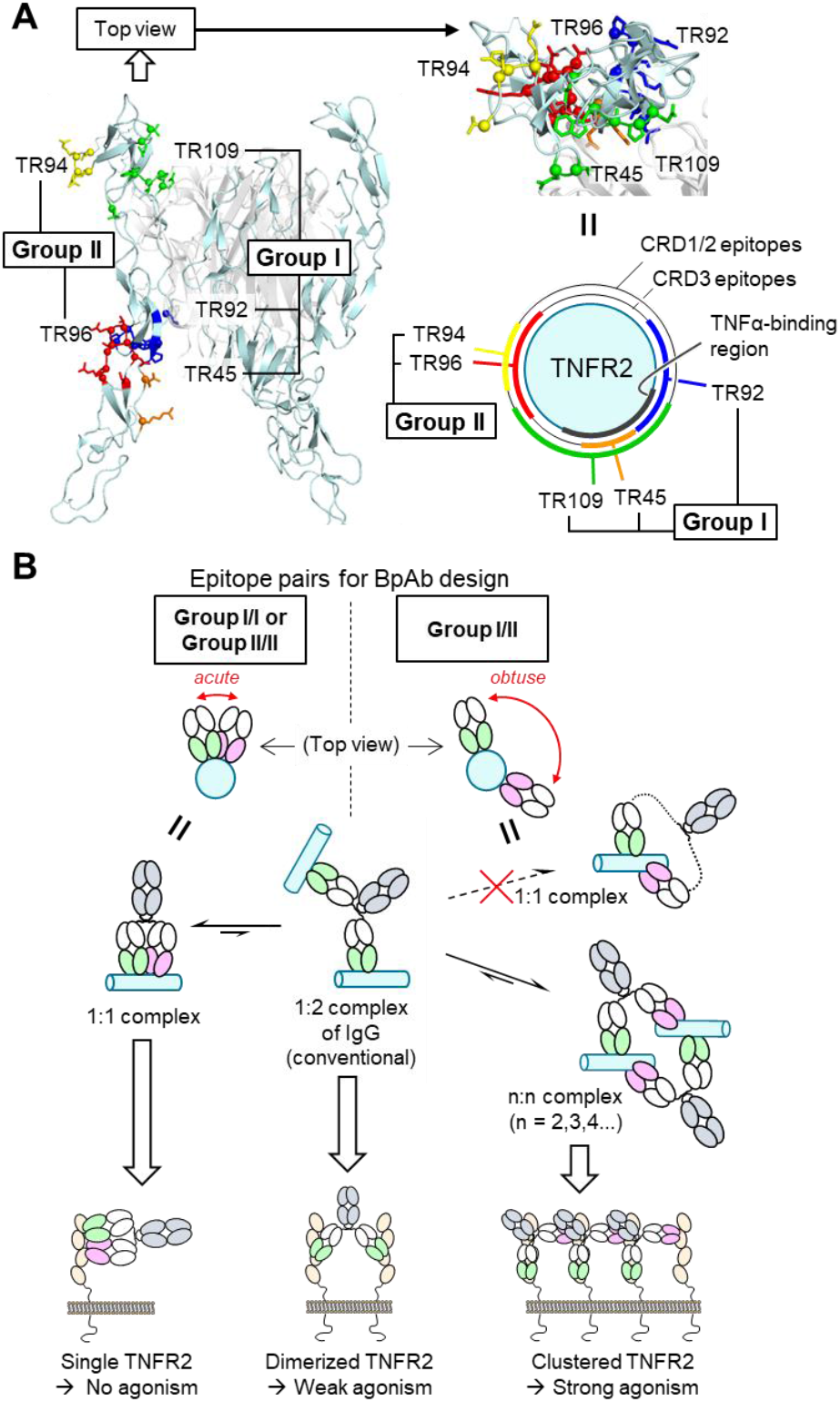
Epitope selection to control agonistic activity. (**A**) Two groups of epitopes for designing biparatopic antibodies (BpAbs). (**B**) Mechanism of control over agonistic activity through crosslinking activity.

In conclusion, we successfully developed various functional BpAbs and elucidated a simple mechanism for regulating the agonistic activity by comprehensively screening bivalent BpAbs against TNFR2. In particular, we developed Bp109-92, which had strong antagonistic activity without any agonistic activity. It was unique as it only had a specific 1:1-binding mode as identified by the cryo-EM structural analysis. The observed Fab-Fab contact may also be advantageous in terms of binding affinity and structural specificity. Although further study is required for investigating if this type of antagonist against TNFR2 can show clinical efficacy, this study serves as a template for designing BpAbs with the desired biological activities. The possible limitation of this design lies in the structure of the target, as 1:1-binding requires a large, exposed surface of the target protein to cover the two epitopes at an appropriate distance. Despite this, we believe that BpAbs with similar design against TNFRSF and other cell-surface receptors would expand the potential of the antibody therapeutics.

## Materials and Methods

### Cell culture

RamosBlue cells expressing human TNFR2 were cultured in IMDM supplemented with 2 mM glutamine and 10% FBS, as previously described (*31*). HEK293T cells were obtained from American Type Culture Collection (ATCC) and were cultured in D-MEM supplemented with 10% FBS. Human peripheral blood mononuclear cells (PBMC) from healthy donors were obtained from the Community Blood Bank (Sioux Falls, SD, USA) and cultured in RPMI1640 supplemented with 2 mM glutamine and 10% FBS.

### Competitive binding analysis

The topographical relationships between the binding of the five anti-TNFR2 mAbs (TR45, TR92, TR94, TR96, and TR109) were analyzed by the mutual competition of all possible pairs of mAbs (5 × 5 = 25) in a label-free competitive ELISA (*32,33*). Each indicator mAb#1 was captured by goat anti-mouse IgG Fc that had been coated on the microtiter plates. In a separate tube, an excess amount of competitor mAb#2 was diluted in blocking buffer, mixed with 20 ng/ml of TNFR2-rabbit Fc fusion protein (*31*), and incubated overnight at 4 °C. The plates were washed twice and the mixtures in the tubes were transferred to each well. Immune complexes captured on the plates were probed with alkaline phosphatase-conjugated goat anti-rabbit IgG. The binding of mAb#1 to the mAb#2-TNFR2-rabbit Fc immune complex was determined as the percentage of binding without mAb#2. No competition (less than 0) was indicated as 0.

### Cloning, expression, and purification of chimeric recombinant antibodies

To produce chimeric antibodies carrying the constant regions of human IgG1 and Igκ, DNA encoding the variable regions of the heavy and light chains were cloned into pFUSE-CHIg-hG1 and pFUSE2-CLIg-hκ (InvivoGen), respectively. Recombinant chimeric antibodies were produced using the Expi293 Expression System (Thermo Fisher Scientific) according to the manufacturer’s protocol. cIgGs were purified from the culture supernatant using a 1 mL HiTrap Protein A HP Column (Cytiva). The expression and purification of biparatopic antibodies followed a previously reported method (*31*). Briefly, Fab **‘N’** (see Fig. 1d) was fused to Cfa Int^N^-maltose binding protein (MBP), and Fab **‘C’** was fused to an Fc with ‘knob’ mutation of knobs-into-holes method, which was co-expressed with an MBP-Cfa Int^C^ fusion of Fc with ‘hole’ mutation to produce a monovalent antibody. The two proteins were mixed in the presence of 2 mM Tris(2-carboxyethyl)phosphine (TCEP) and incubated at 37 °C for 2 h. Disulfide bonds were restored by dialysis against 1 mM oxidized glutathione, and following affinity purification using amylose resin (New England Biolabs) to remove MBP-fused side products, BpAbs were purified in a Superdex 200 Increase 10/300 GL column (Cytiva). The antibodies were stored in phosphate buffered saline (PBS, pH 7.4).

### Preparation of recombinant TNFR2 for physicochemical analysis

Recombinant TNFR2 was obtained from the Fc-fused TNFR2 drug, etanercept (Pfizer, Inc.). IdeS, an Fc-specific protease, was used to isolate the extracellular domains of TNFR2 from etanercept. IdeS with a C-terminal hexahistidine tag in PBS (2 mg/mL) (*34*) was added to solubilized etanercept at 10% (w/w), and the solution was incubated at 37 °C for 1 h. After the reaction, IdeS was removed using a 0.5-mL cOmplete-HisTag Purification Resin column. The column was washed with 2 mL of PBS. The flow-through and wash fractions were then flowed through a 0.5-mL rProtein A Sepharose Fast Flow (Cytiva) column to remove unreacted etanercept and fused Fc. The column was washed with 2 mL of PBS. The flow-through and wash fractions contained the hinge-mediated dimer of extracellular domains of TNFR2, which was further purified in a HiLoad Superdex200 16/600 column. To obtain monomeric TNFR2 extracellular domains, dimeric TNFR2 was reduced using 50 mM 2-mercaptoethylamine at 37 °C for 2 h. The solution was dialyzed in PBS containing 1 mM EDTA for 4 h and then dialyzed in PBS for three days, exchanging the buffer every day. The produced monomeric TNFR2 was purified on a HiLoad Superdex200 16/600 column, and after concentration on an ultrafiltration unit (Amicon-Ultra-15, 10 K), size-exclusion chromatography was conducted using a Superdex 200 Increase 10/300 GL column to remove aggregated proteins.

### Surface plasmon resonance

Interaction of cIgGs and BpAbs with recombinant TNFR2 was monitored using a Biacore T200 instrument (Cytiva). Measurements were performed in PBS supplemented with 0.005% Tween 20 (pH 7.4) at a flow rate of 30 μL/min at 25 °C. The antibodies were captured on a CM5 chip immobilized with anti-human IgG Fc using a Human Antibody Capture Kit (GE Healthcare), according to the manufacturer’s protocol. The antibodies (1 μg/mL) were run for 120 s to capture approximately 400 RU. Analytes were used in a two-fold dilution series of 1.25 – 80 nM (TR94 cIgG), 0.156 – 10 nM (cIgG and BpAbs bearing TR96 variable region), 0.313 – 10 nM (Bp94-92 and Bp109-92), 0.625 – 20 nM (TR92 cIgG), or 0.625 – 40 nM (others). The contact and dissociation times for the antigen were 90 and 300 s, respectively, and the anti-human IgG Fc antibody was regenerated by 3 M MgCl_2_ run for 30 s. The kinetic parameters were obtained by 1:1 global kinetics fitting using Biacore T200 Evaluation Software.

### Flow cytometry

The cells were analyzed using a BD LSRFortessa Cell Analyzer (BD Biosciences). The immunochemical reagents used were as follows: anti-human IgG, Fcγ-PE (Jackson Immunoresearch, #109-116-170), CD3-BV711 (BioLegend, #300463), CD4-BV510 (BioLegend, #344633), CD25-BV421 (BioLegend, #302629); CD127-PerCP-Cy5.5 (BD, #560551), TNFR2-PE (R&D Systems, #22235), and Foxp3-AlexaFluor 647 (BioLegend, #320313). Cells were treated with PBS containing 0.2% sodium azide and 5% FBS. For antibody-binding analysis of TNFR2-expressing RamosBlue and HEK293T cells, primary antibodies were first incubated in a dilution series for 30 min on ice and then labeled with anti-human IgG-PE. The data were analyzed using FlowJo_v10.8.1 (FlowJo, LLC).

### Epitope mapping

The epitopes of the five mAbs were determined using the multiple constructs listed in Table S1. Vectors encoding human TNFR2, mouse TNFR2, and partially substituted mutants were constructed in an expression vector based on pcDNA3.1, which contains an internal ribosome entry site element to co-express TNFR2 with a reporter TagBFP. The expression vector was transfected into HEK293T cells using PEI’MAX’ (Polysciences Inc.). Cells were cultured for 40 h post-transfection and detached from the culture vessels using trypsin/EDTA for 5 min. Cells expressing wild-type and mutant TNFR2 were labeled covalently by combinations of succinimidyl ester compounds of Pacific Orange, DyLight 633, or DyLight 800 (Cat Nos. P30253, 46421, and 46414; Thermo Fisher Scientific). The labeling agents were mixed in 12 different combinations of Pacific Orange (1 mg/mL), DyLight 633 (0.03 mg/mL), DyLight 800 (0.01 mg/mL or 0.3 mg/mL) or their absence, and the cells were incubated with the agents in PBS, pH 7.8 with 3% DMSO for 20 min at 31 °C. Twelve differently labeled cells were mixed, mAbs (1.5 μg/mL) and the secondary antibody were incubated, and the cells were analyzed as described above.

### Reporter assay

TNFR2-expressing RamosBlue cells were seeded at 5 × 10^4^ cells/well in 100 μL of medium containing antibodies at the indicated concentrations in the absence or presence of 50 ng/mL TNFα (Peprotech #AF-200-01A). The cells were incubated for 18 h, and secreted alkaline phosphatase was analyzed using p-nitrophenyl phosphate. Colorimetric changes were detected by measuring absorbance at 405 nm. Signals were normalized to the average of eight wells incubated without antibody or TNFα (negative) and eight wells incubated only with 50 ng/mL TNFα (positive). For the chart showing the agonistic activity, the negative and positive values were set at 0.1 and 0.9, respectively. For charts showing antagonistic activity, the values were set at −0.9 and −0.1, respectively.

### Size determination of immunocomplexes

Size-exclusion chromatography with a multi-angle light scattering detector (SEC-MALS) was conducted using a Superose 6 Increase 10/300 GL column (Cytiva) and DAWN 6 (Wyatt) as the column and detector, respectively. PBS was used as the running buffer and run at 0.5 mL/min. Under typical conditions, 2 μM antibody was mixed with 2 or 16 μM TNFR2 and incubated for 10–20 min at room temperature before running. Data were analyzed using ASTRA 6 software (Wyatt). The concentration of proteins was calculated from the refractive index using d*n*/d*c* = 0.169. Molar mass values were determined by Debye fitting of angle-dependent light scattering.

Mass photometry was conducted using a Refeyn One (Refeyn Ltd.). Five-microliter sample of BpAbs (50 nM) mixed with TNFR2 (50 or 400 nM) in PBS was diluted four-fold with 15 μL PBS pre-loaded for hydration on a glass slide. Interferometric scattering was observed under a microscope for 1–2 min, and the accumulated interference signal was analyzed as previously described (*21*).

### Negatively-stained electron microscopy

Bp109-92 and Bp45-94 in complex with TNFR2 were prepared in a 1:1 mixture (1.3 μM) and stained without purification. Bp109-92 F(ab′)_2_ was prepared by incubating Bp109-92 before purification with IdeS (1/10 equivalent to Bp109-92) at 37 °C for 2 h. IdeS and Fc were removed using the same method as that used to produce recombinant TNFR2. Bp109-92 F(ab′)_2_ was purified on a Superdex 200 Increase 10/300 GL column. Complex of Bp109-92 F(ab′)_2_ with TNFR2 was prepared as a 1:4 mixture (2.6 μM of F(ab′)_2_ and 10.2 μM of TNFR2) for 30 min at room temperature, and purified in a Superose 6 Increase 10/300 GL column. The sample solutions were diluted to ∼0.1 mg/mL and applied to carbon-coated copper grids that had been glow-discharged for 20 s at 20 mA using an ion coater IB-3 (eiko) and negatively stained with 2% (w/v) uranyl acetate. Micrographs were recorded at a nominal magnification of 25,000×, corresponding to 8 Å/pixel, using a JEM-1011 transmission electron microscope (JEOL, Tokyo, Japan) operating at 100 kV with a TemCamF416A-Hs-4 CMOS camera (TVIPS). Image processing procedures, including particle selection, 2D classification, and averaging were performed using the RELION program (*36*).

### Cryo-EM specimen preparation and data collection

The maltose-binding protein fused to the four cysteine-rich domains of TNFR2 (TNFR2CRD-MBP; sequence in Fig. S13) was cloned into pcDNA3.1. Protein expression was determined using the Expi293 Expression System (Thermo Fisher Scientific). Six days post-transfection, the culture supernatant was dialyzed overnight against Buffer A (20 mM Tris, 300 mM NaCl, pH 8.0) containing 5 mM imidazole. Expressed proteins were captured on Ni-NTA Superflow (Qiagen) equilibrated with Buffer A containing 5 mM imidazole. The resins were washed sequentially using Buffer A containing 5, 10, and 20 mM imidazole, and the protein was eluted using Buffer A containing 200 mM imidazole. The eluate was dialyzed against PBS and the final purification was conducted using a Superdex200 16/600 column.

Complex of Bp109-92 and TNFR2CRD-MBP was prepared as a 2:3 mixture (5.6 μM Bp109-92 and 8.4 μM TNFR2CRD-MBP) for 5 min at room temperature, and purified in a Superose 6 Increase 10/300 GL column. Three microliters of the complex solution (0.2 mg/mL) was applied to a Quantifoil R1.2/1.3 Cu 200 mesh grid (Microtools GmbH) that was glow-discharged for 20 s at 20 mA using a JEC-3000FC sputter coater (JEOL). The grid was blotted with a force of 0 and a time of 3 s in a Vitrobot Mark IV chamber (Thermo Fisher Scientific) equilibrated at 4 °C and 100% humidity, and then immediately plunged into liquid ethane. Excess ethane was removed with filter paper, and the grids were stored in liquid nitrogen. The image dataset was collected using SerialEM (*37*) yoneoLocr (*38*) and JEM-Z300FSC (CRYO ARM 300: JEOL) operated at 300 kV with a K3 direct electron detector (Gatan, Inc.) in CDS mode. The Ω-type in-column energy filter was operated with a slit width of 20 eV for zero-loss imaging. The nominal magnification was 60,000×. Defocus varied between –0.5 and –2.0 μm. Each movie was fractionated into 60 frames (0.081 s each, total exposure:4.87 s), with a total dose of 60 e^−^/Å^2^.

### Cryo-EM image processing and model building

A gain reference image was prepared with the relion_estimate_gain command in RELION 4.0 (*36*) using the first 500 movies. Images were processed using cryoSPARC ver. 3.3.2 (*39*). A total of 5,724 movies were imported, motion corrected, and contrast transfer functions (CTFs) were estimated. A total of 4,160 micrographs with maximum CTF resolutions greater than 5 Å were selected. First, the particles were automatically picked using a blob picker job with a particle diameter between 100 and 150 Å. After particle extraction with 4x binning, 2D classification into 50 classes was performed to select clear 2D class averages as templates for subsequent particle picking. Then, the particles were automatically picked again with the templates, and 2,218,071 particle images were extracted with a box size of 320 pixels using 4x binning (downscaled to 80 pixels). Two rounds of 2D classification into 50 classes, with a circular mask of 170 Å, were performed to select 354,477 particles. The number of final full iterations and the batch size per class were increased to 20 and 200, respectively. The best initial model was selected from four reconstructed models. A total of 100,391 particles belonging to the best model were extracted again with a box size of 320 pixels without binning. After homogeneous refinement, global/local CTF refinement and nonuniform refinement (*40*) were performed to reach 4.26 Å overall map resolution. The particle images were downsampled to 256 pixels (corresponding to a pixel size of 1.088 Å). After one round of non-uniform refinement, the handedness of the map and mask was flipped. After another round of non-uniform refinement and two rounds of local CTF refinement and non-uniform refinement, a final map was reconstructed at 3.63 Å resolution (FSC=0.143).

Homology models of Fab109 and Fab92 were generated using SWISS-MODEL (*41*). The atomic model of TNFR2 (PDB entry:3ALQ) and the homology models of Fabs were manually fitted into the density and modified using UCSF Chimera (*42*) and Coot (*43*). Real space refinement was performed using the PHENIX software (*44*). Several rounds of manual model modification and real space refinement were repeated. Figures were prepared using UCSF Chimera (*42*) and PyMOL (Schrödinger, LLC). The parameters are summarized in Table S3.

### Stimulation of PBMCs

PBMCs were passed through a 40-μm cell strainer and diluted to 1 × 10^5^ cells/well in a 96-well round-bottom plate, and were cultured in the presence of 10 ng/mL IL-2 (Genscript #Z00368), and the presence or absence of TNFα (100 ng/mL) and antagonists (50, 150, and 500 ng/mL). The cells were incubated for 48 h and analyzed by flow cytometry. For flow cytometry, dead cells were labeled using the LIVE/DEAD™ Fixable Near IR (780) Viability Kit (Thermo Fisher Scientific). eBioscience Intracellular Fixation & Permeabilization Buffer Set (Thermo Fisher Scientific) was used to stain Foxp3. BD CompBeads (BD) were used to compensate for the fluorophores. One thousand steps of iteration by fast Fourier transform-accelerated interpolation-based t-distributed stochastic neighbor embedding (FIt-SNE) implemented in FlowJo v10.8.1, was conducted for cluster analysis (*45*).

## Acknowledgments

The authors thank Reiko Satoh and Mayumi Niiyama for their technical assistance with protein production, Dr. Yasuhiro Abe, Akiko Abe, Takahide Mori, Miho Mukai, and Sayuri Okamoto for their assistance with the production of antibody panels, and Dr. Kohei Shiba for assistance with mass photometry.

## Author contributions

Conceptualization: HA, HK, SN, KT; Investigation: HA, TI, JF, KNi, TM, TK, SN; Funding acquisition: HA, JF, KNa, HK; Writing – original draft: HA, JF, SN; Writing – review and editing: TK, KNa, HO, HK, KT.

## Data and materials availability

Cryo-EM density map and model of Bp109-92 in complex with TNFR2-MBP are deposited to Electron microscopy Data Bank (EMDB) and Protein Data Bank (PDB) with accession codes of EMD-34871 and PDB-8HLB, respectively. All other data are available in the main text or supplementary materials.

