## Supplementary Materials for "Development of a 1:1-binding biparatopic anti-TNFR2 antagonist by epitope selection"

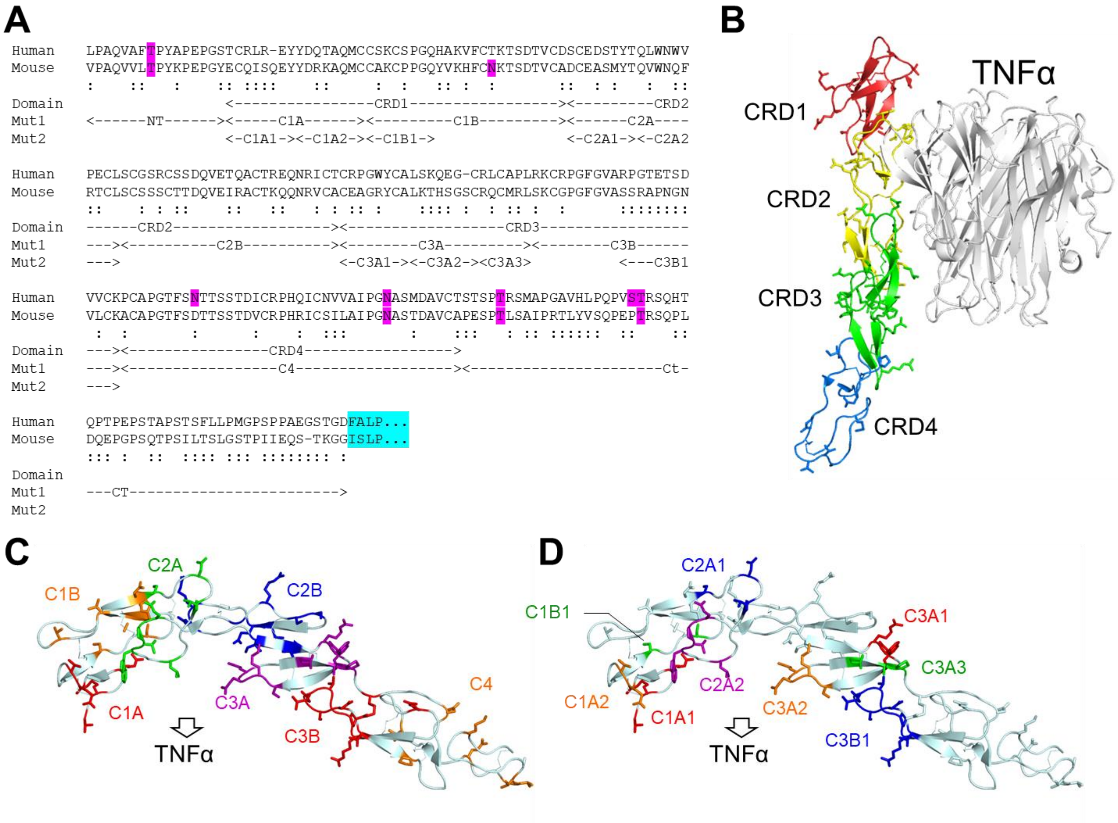


**Fig. S1. Design of mutants for determining the antibody epitopes. (A)** Alignment of the extracellular regions of human TNFR2 (UniProt ID: P20333) and mouse TNFR2 (UniProt ID: P25119) and the analyzed mutants. The residues colored purple are the potential glycosylation sites and the residues colored cyan are the N-termini of transmembrane helix. ':' indicates different amino acids between the two orthologs. Human-to-mouse mutants were designed so that the peptide sequences of the indicated regions on Mut1 or Mut2 lines were replaced to that of mouse TNFR2. Mouse-to-human mutants were designed in the opposite way. **(B)** Domains mapped onto the tertiary structure of human TNFR2 in complex with TNFα (PDB ID: 3ALQ). One TNFR2 molecule and trimeric TNFα are shown. **(C,D)** Mutated amino acids of each mutant in the Mut1 (**C**) and Mut2 (**D**) series are labeled on the structure of TNFR2. In **B**–**D**, CT and NT regions are disordered and not shown.

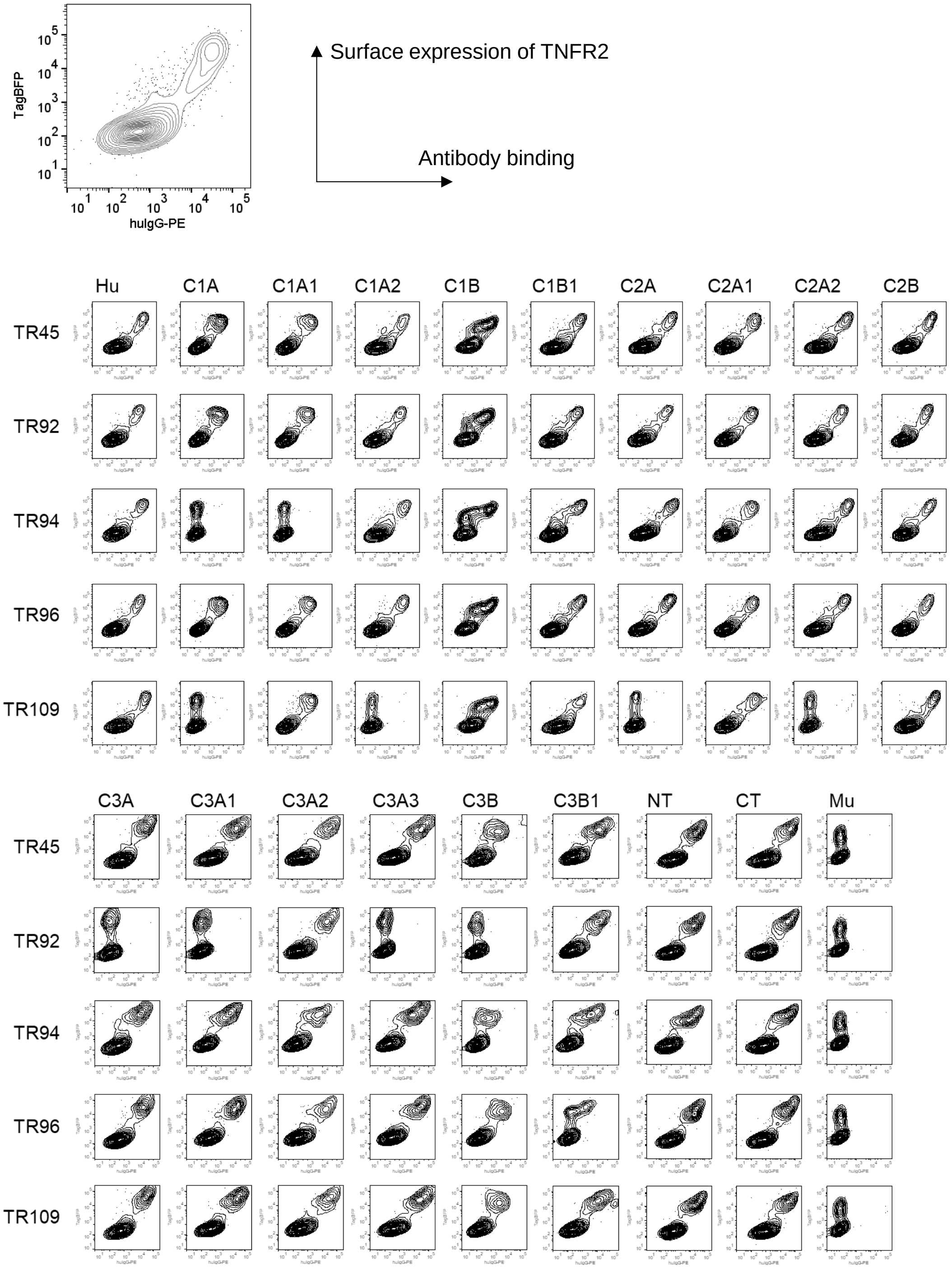


**Fig. S2. Binding of monoclonal antibodies to the wild-type and mutant human TNFR2 series.** Expression of TNFR2 was visualized by TagBFP as a bicistronic reporter (y-axis). Binding of the antibody was visualized by secondary antibody conjugated with R-PE (x-axis). Reduced antibody binding to wild-type or human-to-mouse mutant TNFR2$^{+}$ cells indicates that the mutated region is the epitope of the antibody.

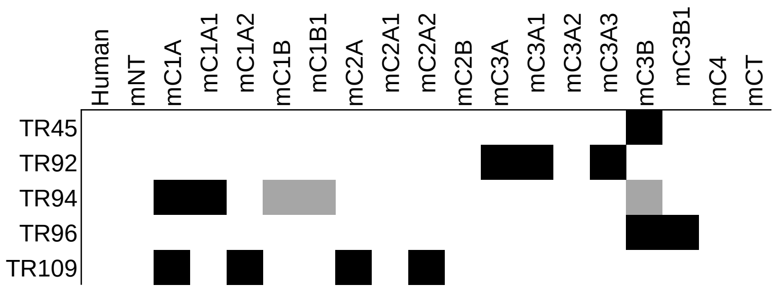


**Fig. S3. Simplified map of epitope regions determined by binding reduction.** The regions were determined by reduced binding of the monoclonal antibodies to the cells expressing wild-type human TNFR2 or the mutants with the indicated regions substituted by peptides from mouse TNFR2. Black indicates loss of binding; grey indicates partially reduced binding. Reduction is analyzed by flow cytometry presented in Fig. S2.

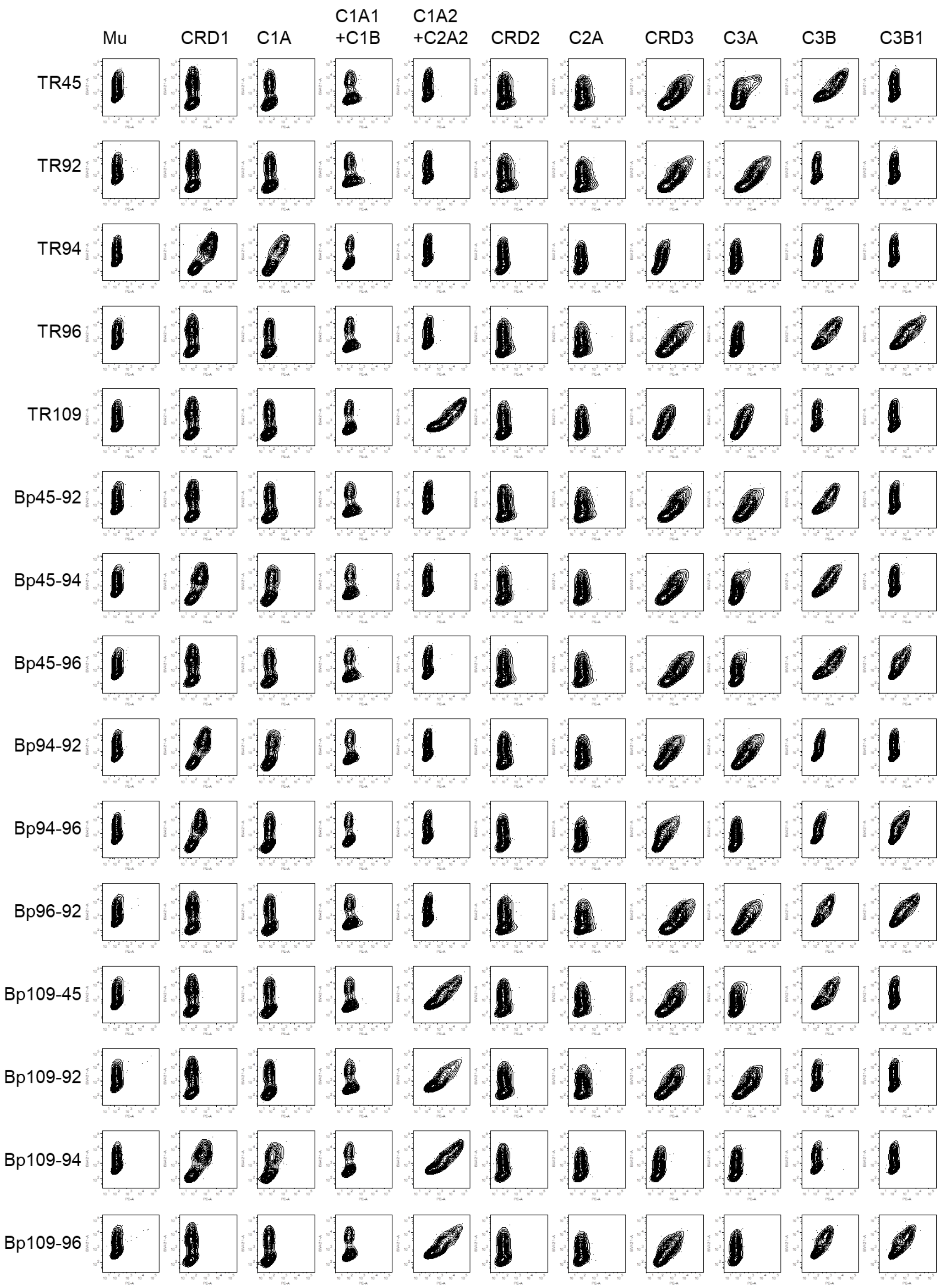


**Fig. S4.** *Caption to the next page.*

**Fig. S4. Binding of chimeric monoclonal antibodies and biparatopic antibodies to the wild-type and mutant murine TNFR2 series.** Expression of TNFR2 was visualized by TagBFP as a bicistronic reporter (y-axis). Binding of the antibody was visualized by secondary antibody conjugated with R-PE (x-axis). Antibody binding to mouse-to-human mutant TNFR2$^+$ cells indicates that the mutated region is the epitope of the antibody.

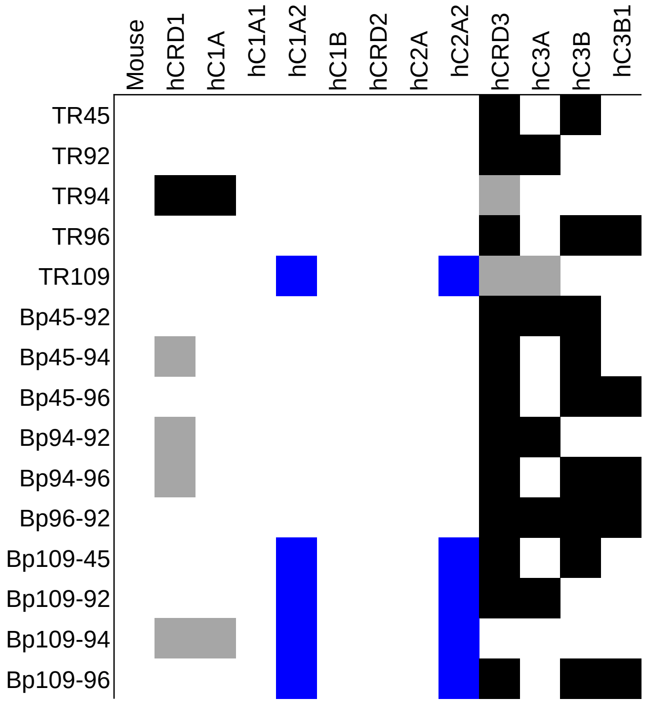


**Fig. S5. Simplified map of epitope regions determined by binding.** The regions were determined by increased binding of the monoclonal antibodies to the cells expressing wild-type mouse TNFR2 or the mutants with the indicated regions substituted by peptides from human TNFR2. Black indicates gain of binding; grey indicates partially gained binding;blue indicates gain of binding when two regions were mutated at the same time. Reduction is analyzed by flow cytometry presented in Fig. S4.

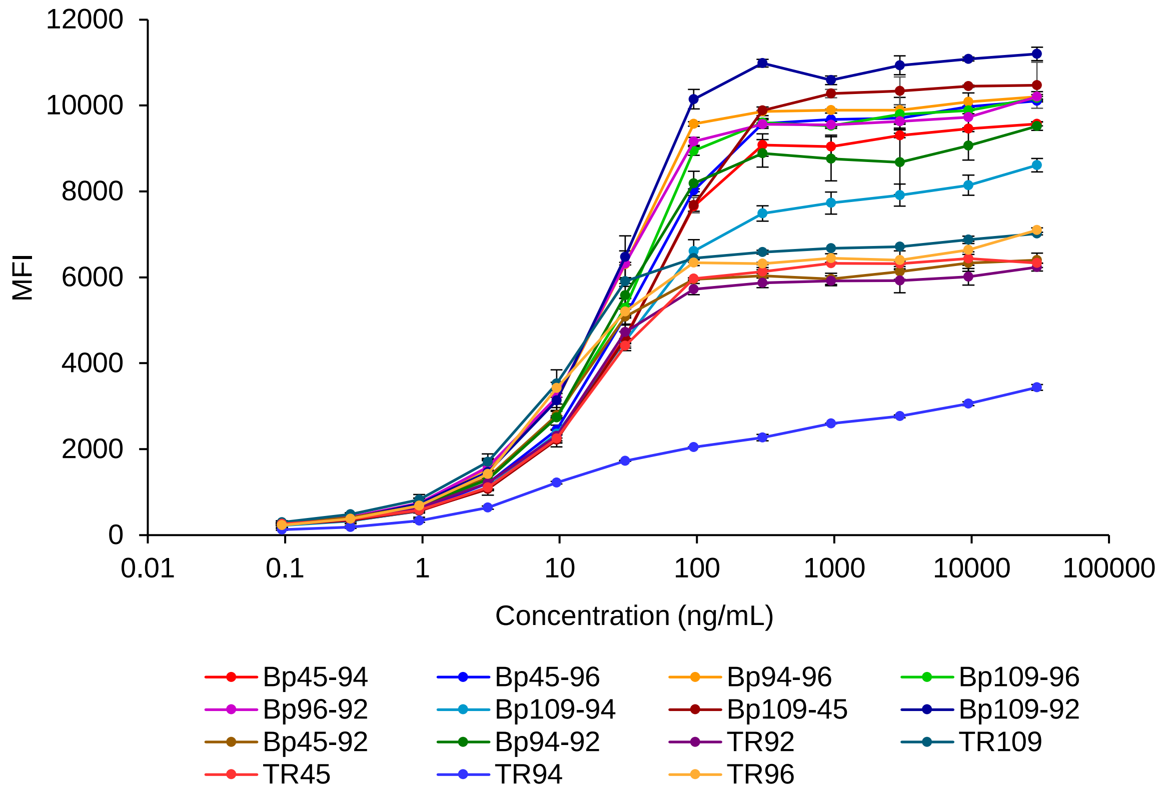


**Fig. S6. Concentration-dependent binding of antibodies to TNFR2-expressing RamosBlue cells.** Binding was detected using anti-human IgG R-PE, and mean fluorescence intensity (MFI) was obtained in two independent experiments. To standardize between the two experiments, the MFI values for three highest concentrations were averaged for each antibody (= $A_i$; i, each antibody). $\Sigma(kA_i^{1st} - A_i^{2nd})$ was minimized for $A$ values of ten antibodies excluding those with TR94 variable regions as they did not reach a plateau. Using the standardized MFI, the average values are shown with standard deviation.

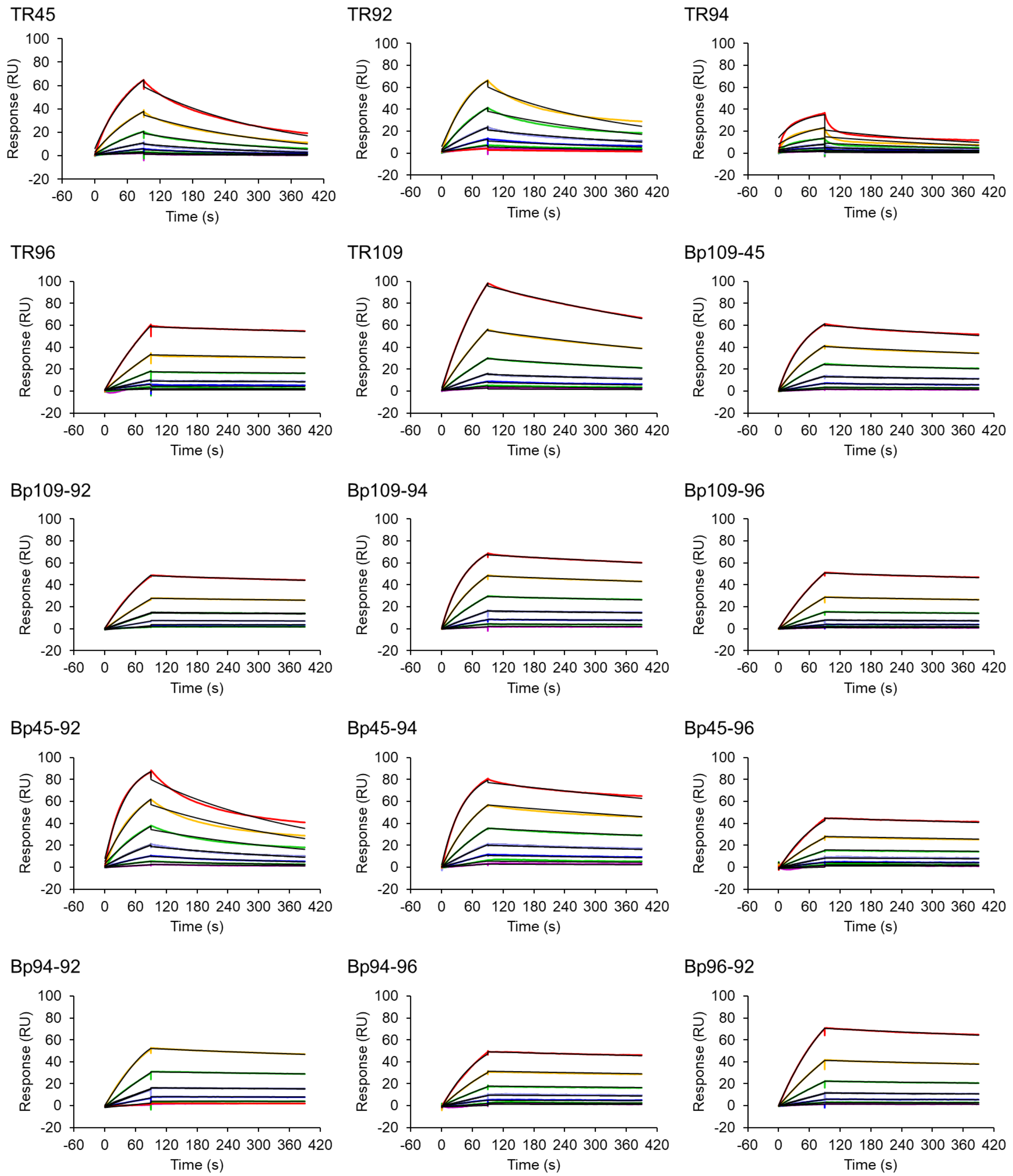


**Fig. S7. Surface plasmon resonance sensorgrams of the antibody binding to TNFR2.** See Materials and Methods for the concentration analyzed and Table S2 for the kinetic parameters. Binding of TR94 was not strong and the parameters were determined poorly by fitting to 1:1 binding kinetics.

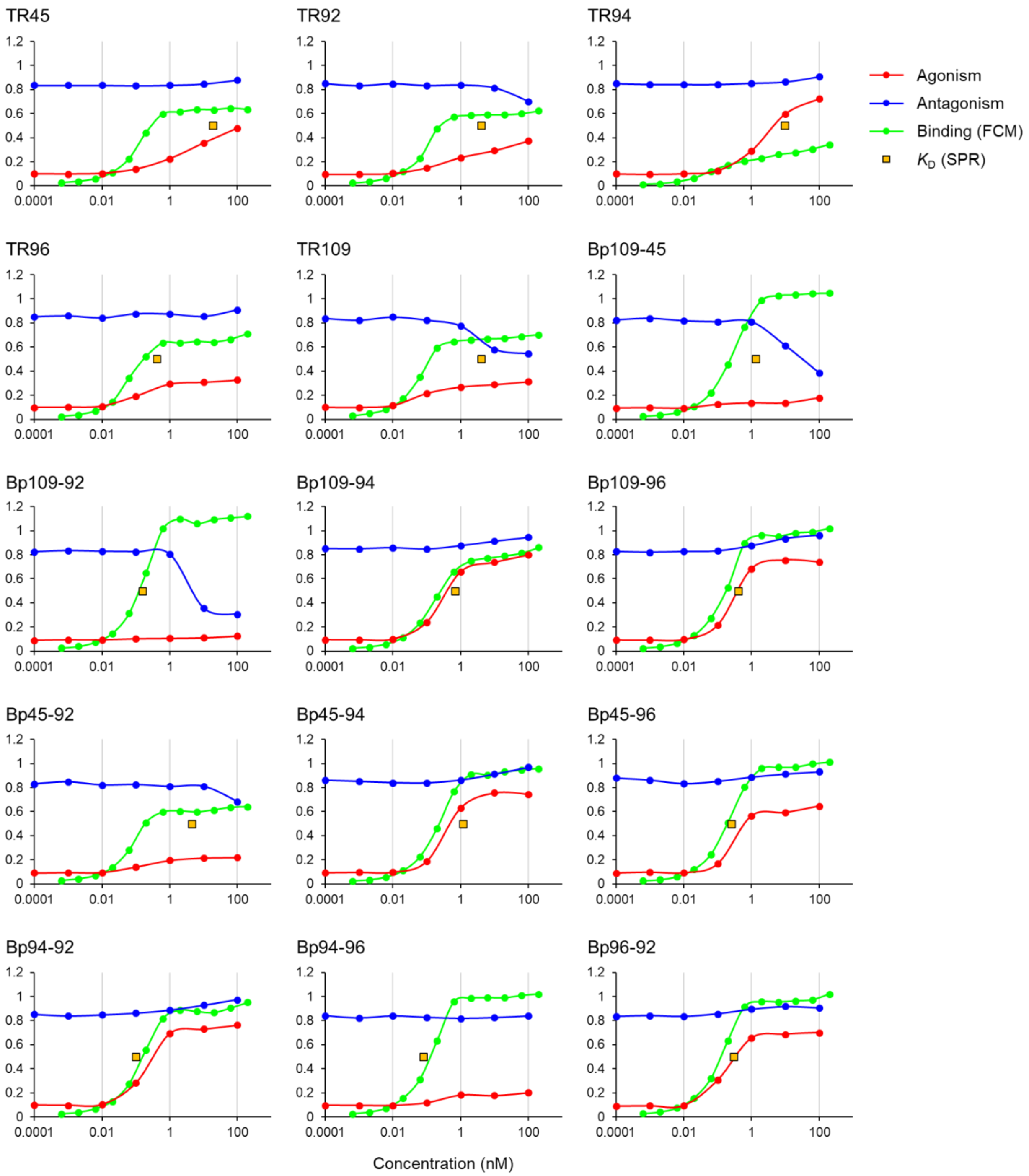


**Fig. S8. Comparative analysis of biological and binding activities of the antibodies.** For all panels, x-axes are concentration (nM) and y-axes are arbitrary units shared for all 15 antibodies. Data are adopted from Fig. 2E, Fig. S7, and Table S2. Red, biological activity (upregulation by agonistic activity); blue, biological activity in the presence of TNFα (downregulation by antagonistic activity); green, binding activity by flow cytometry; yellow square, dissociation constant against TNFR2.

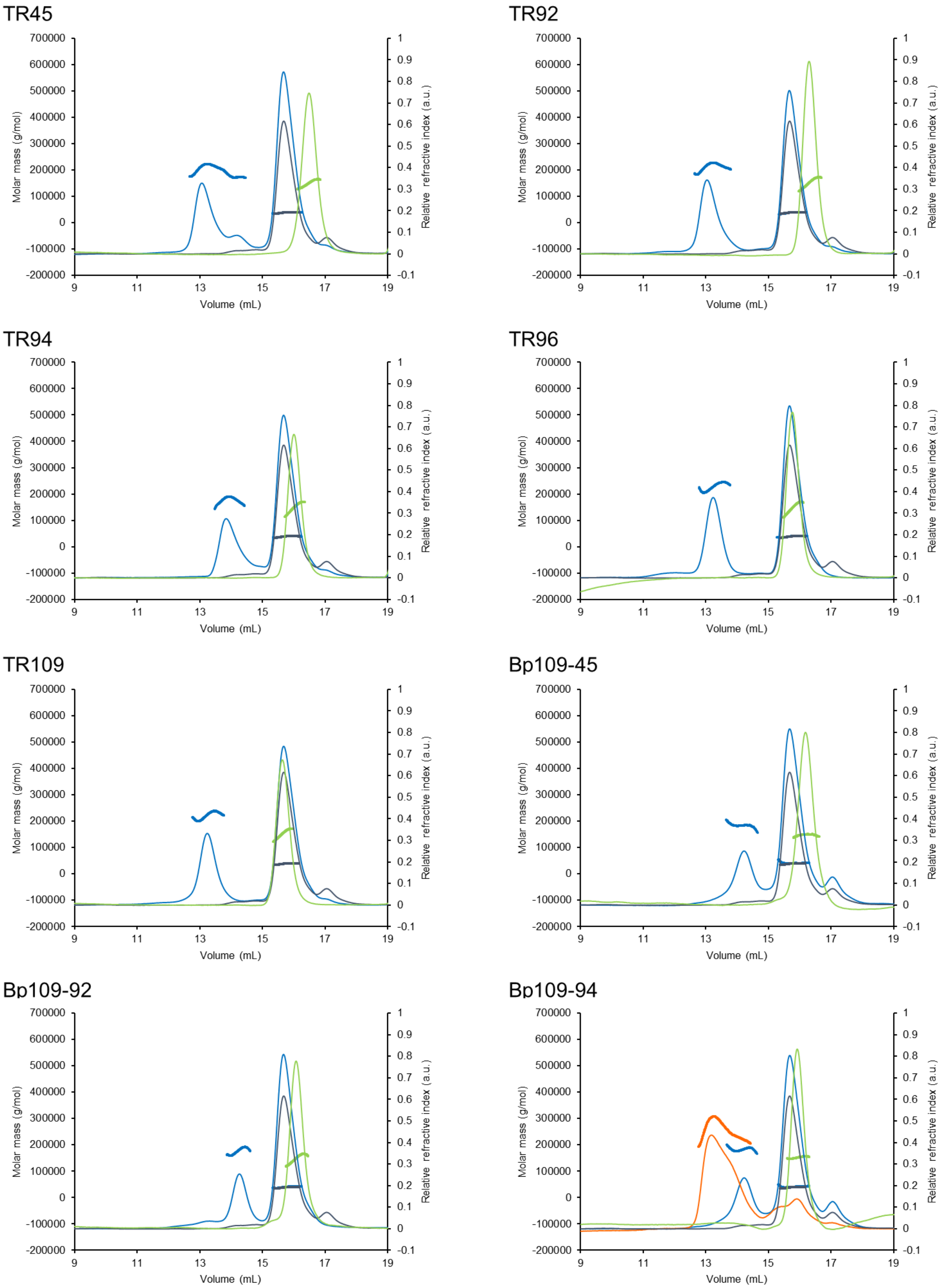


**Fig. S9.** *Continued to next page.*

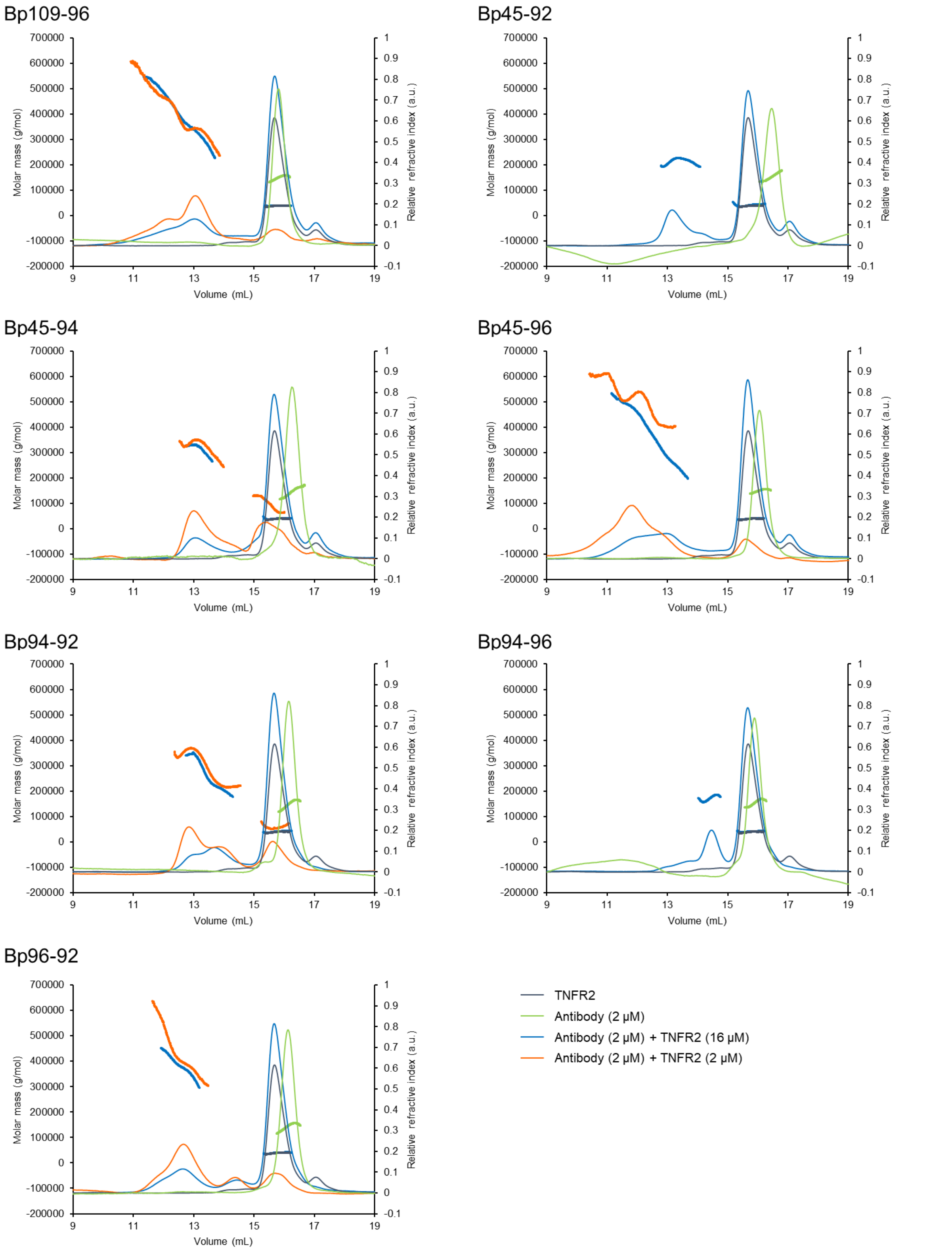


**Fig. S9. SEC-MALS charts.** For each chromatogram, relative refractive index is shown with thin lines (right y-axis), and the molar mass is shown with large dots (left y-axis).

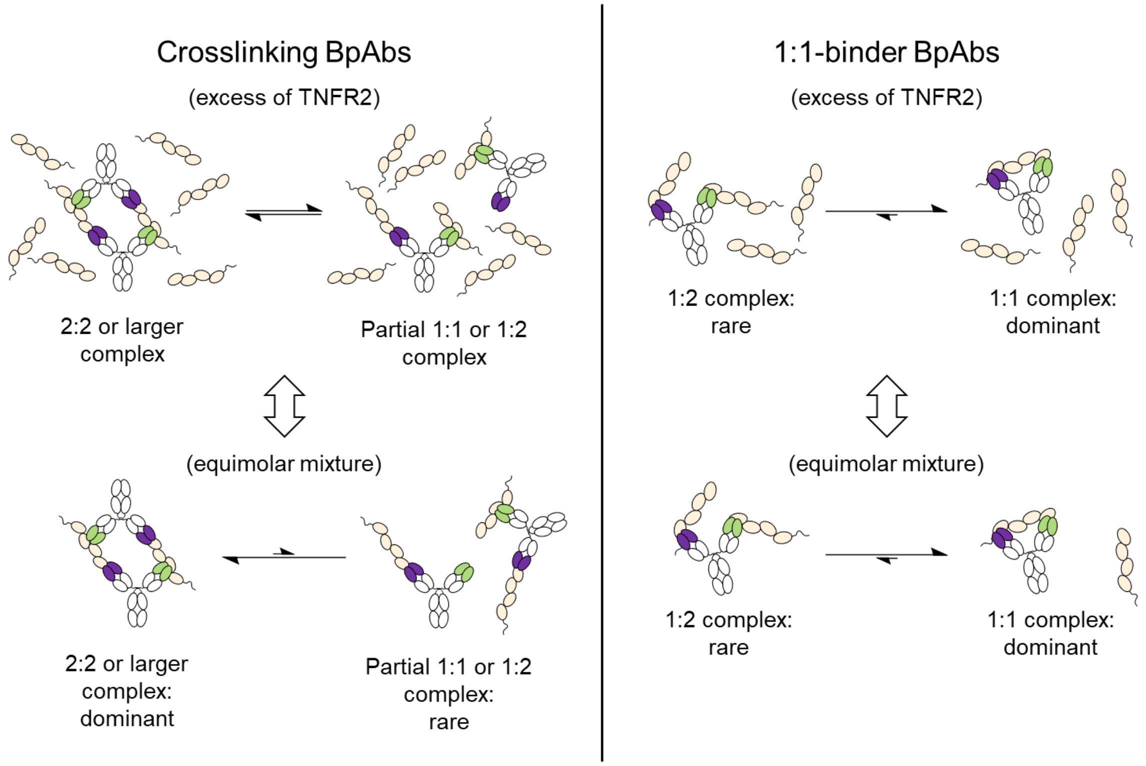


**Fig. S10. Proposed equilibrium of complex formation in solution using recombinant TNFR2.** In the case of bridging BpAbs (left), large complexes and partial 1:1 or 1:2 complexes are in equilibrium in the presence of excess TNFR2. On the other hand, a large complex is dominant for equimolar mixture due to the absence of sufficient amounts of TNFR2 for 1:2 complex formation (Fig. 3E,F). In case of 1:1-binder BpAbs, 1:1 complex is dominant irrespective of the BpAb:TNFR2 ratio (Fig. 3D).

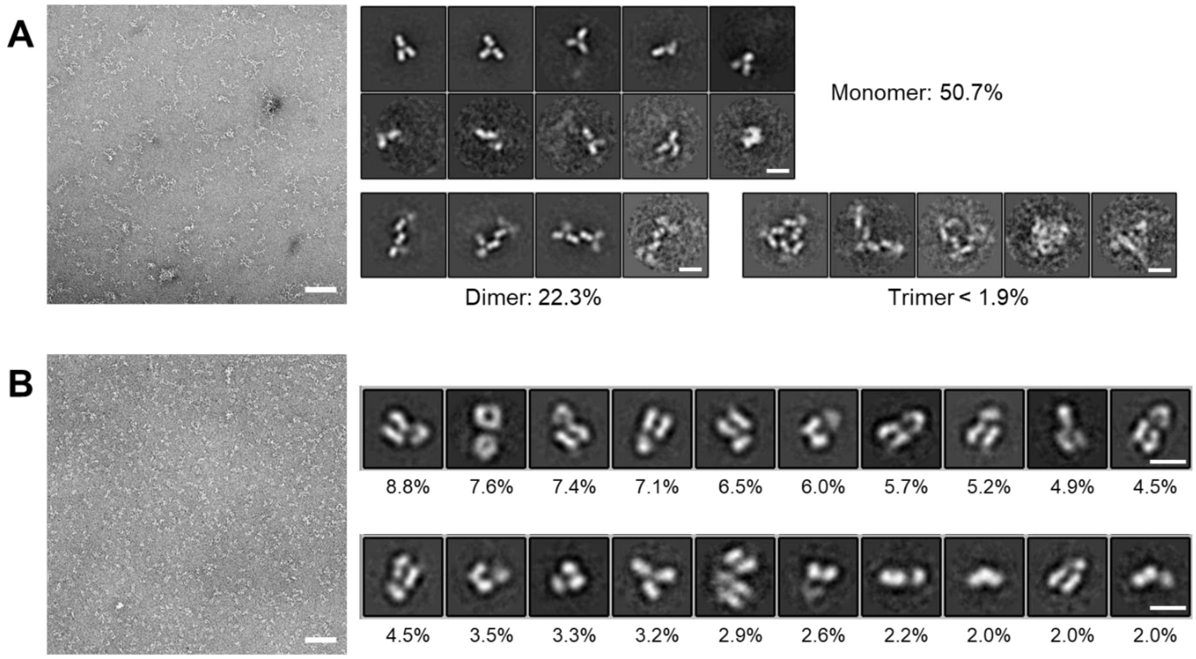


**Fig. S11. Negatively stained electron microscopic images of selected biparatopic antibodies in complex with TNFR2.** A) Bp45-92, B) Bp109-92. Left, representative images (Scale bar: 500 Å); right, particles classified and the percentage of each class of structures in 6,725 (A) and 9,786 (B) particles (Scale bar: 100 Å). For A, classes are defined by the number of IgG-like structural elements per particle. For B, top 20 classes are shown.

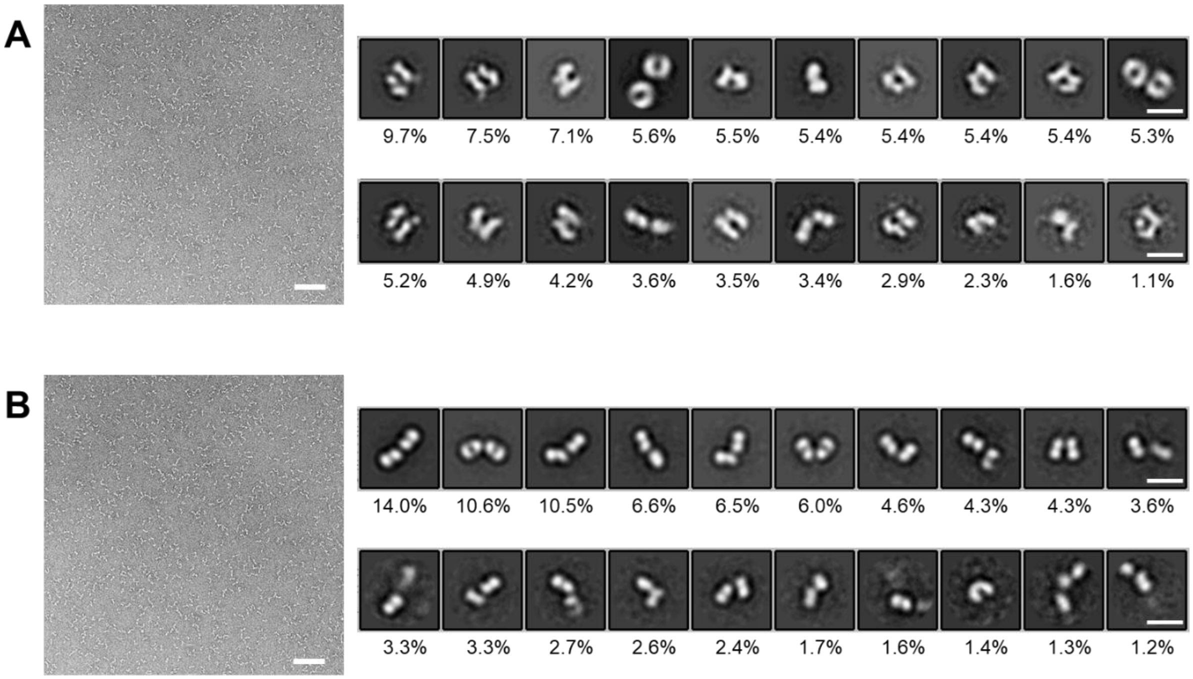


**Fig. S12. Negatively stained electron microscopic images of Bp109-92 F(ab′)$_2$ in the A) presence or B) absence of TNFR2.** Left, representative images (Scale bar: 500 Å); right, particles classified and the percentage of each class of structures in 12,267 (A) and 13,046 (B) particles (Scale bar: 100 Å). Top 20 classes are shown.

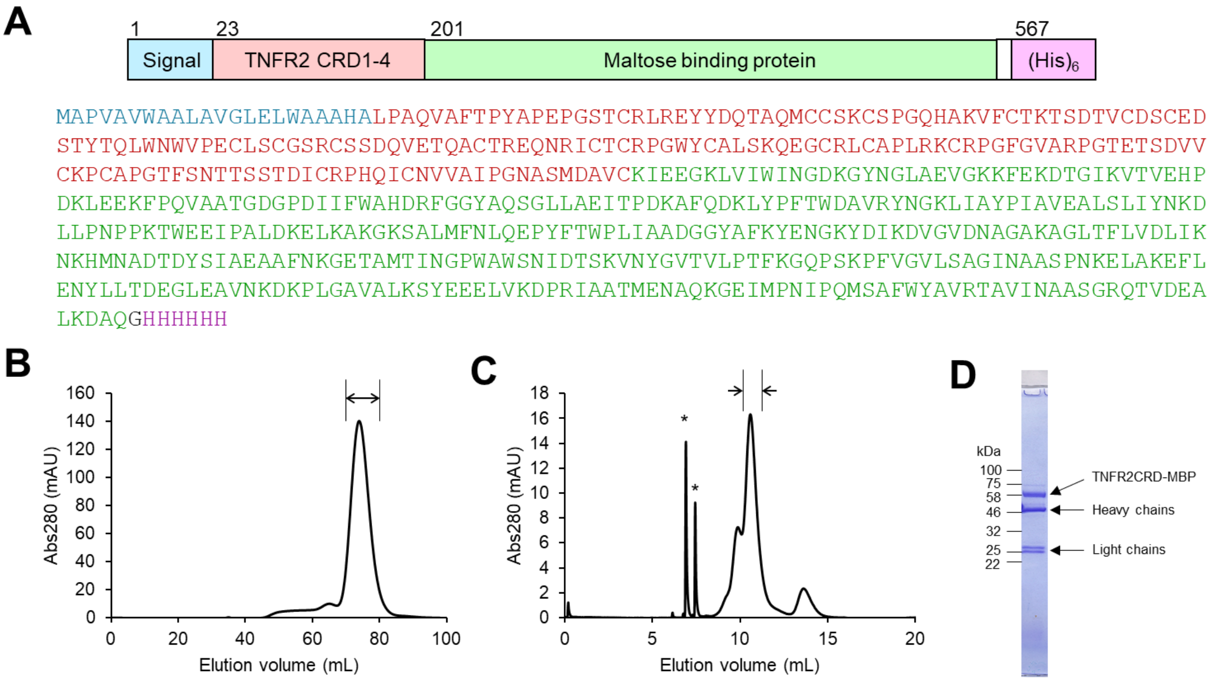


**Fig. S13. Preparation of the samples for cryo-electron microscopy.** (**A**) The TNFR2 construct used. (**B**) Size-exclusion chromatogram of TNFR2CRD-MBP using Superdex200 16/600 column. The arrow indicates the collected fractions. (**C**) Size-exclusion chromatogram of Bp109-92–TNFR2-MBP complex using Superose6 Increase 10/300 column. The arrows indicate the collected fractions. Asterisks are unrelated signals due to accidental stop of the fraction collector. (**D**) CBB-stained SDS-PAGE of the obtained complex.

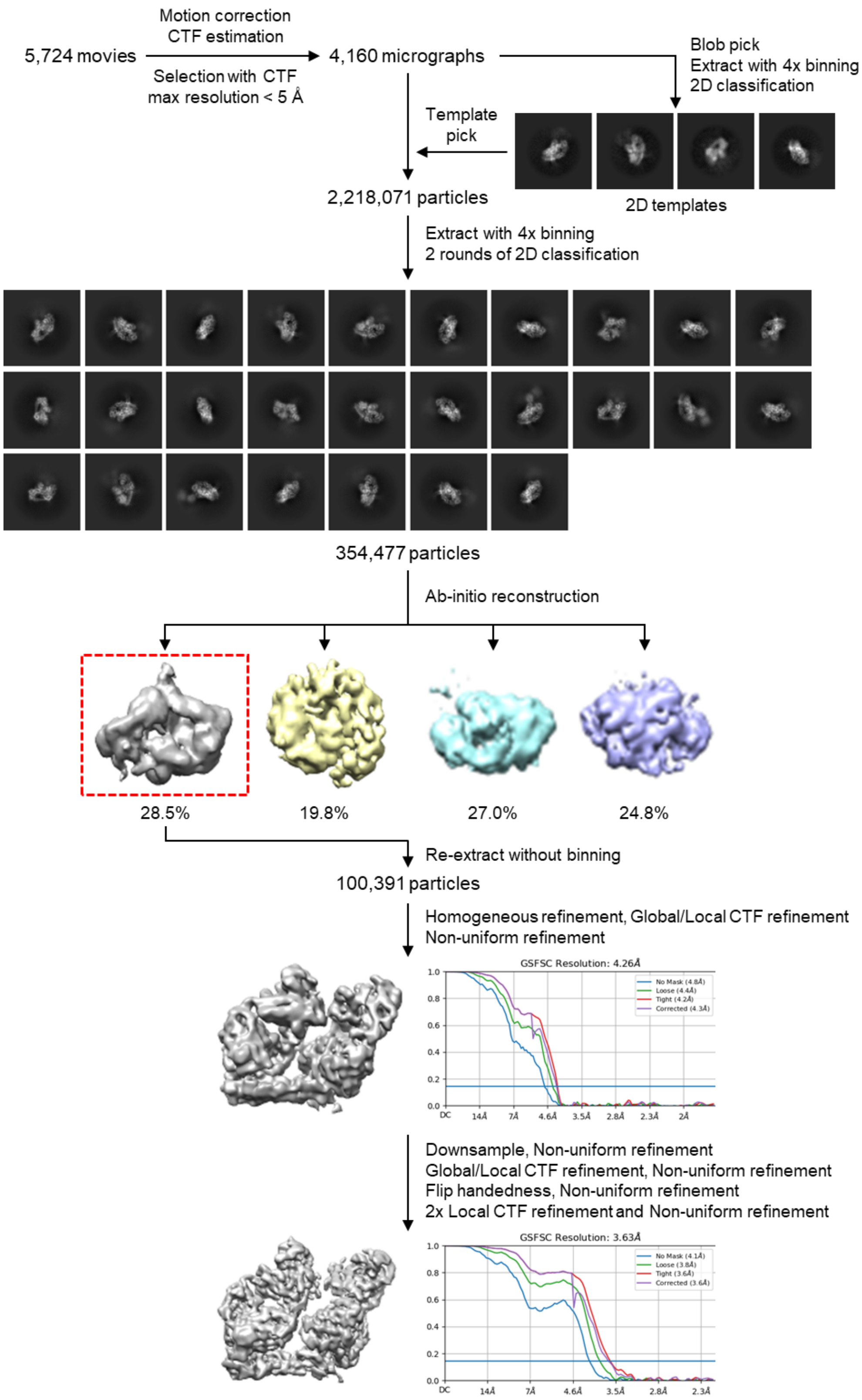


**Fig. S14. Image processing of Bp109-92–TNFR2-MBP complex.**

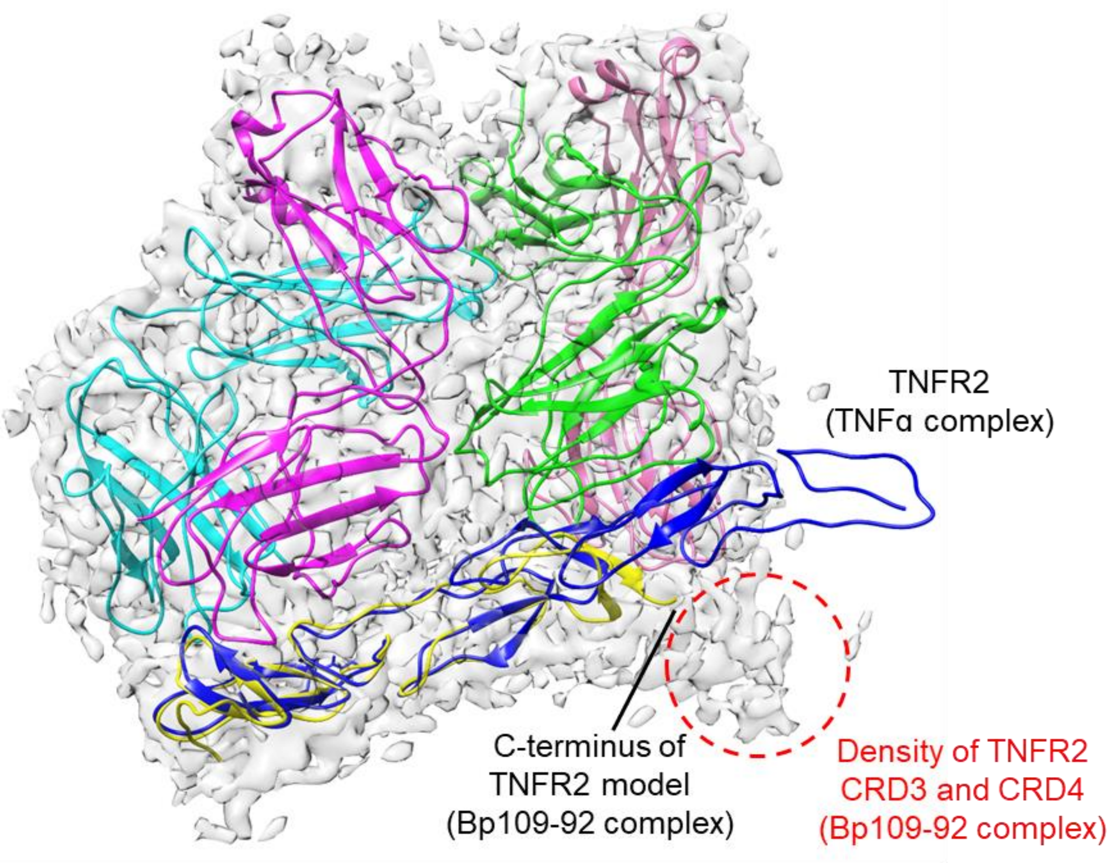


**Fig. S15. Structure comparison of Bp109-92–TNFR2-MBP complex with TNFα–TNFR2 complex.** Overall view of the cryo-EM structure colored as in Fig. 4A. The map is contoured at a lower level than Fig. 4A. TNFR2 in complex with TNFα (blue) (PDB ID: 3ALQ) is superposed with TNFR2 in complex with Bp109-92 (yellow).

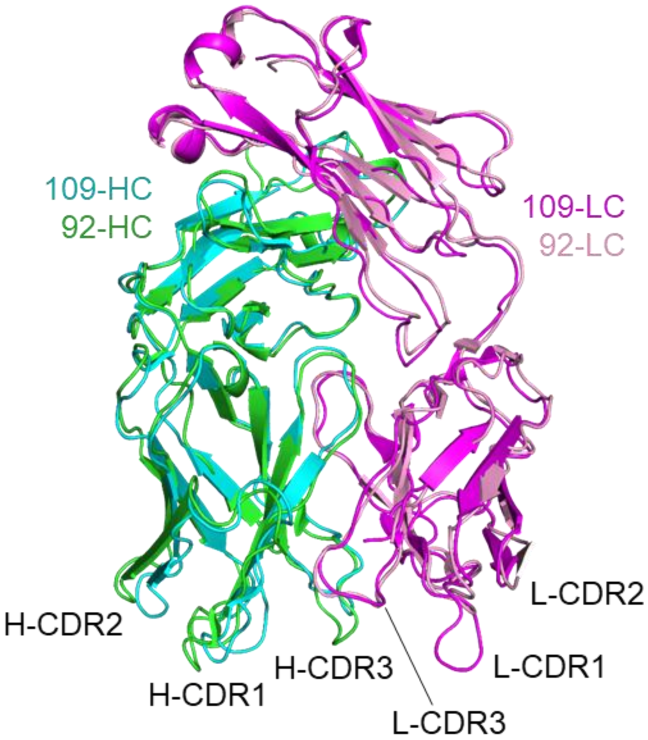


**Fig. S16. Superposition between 109-Fab and 92-Fab in the Bp109-92–TNFR2 complex.** Chains are colored as in Fig, 4A.

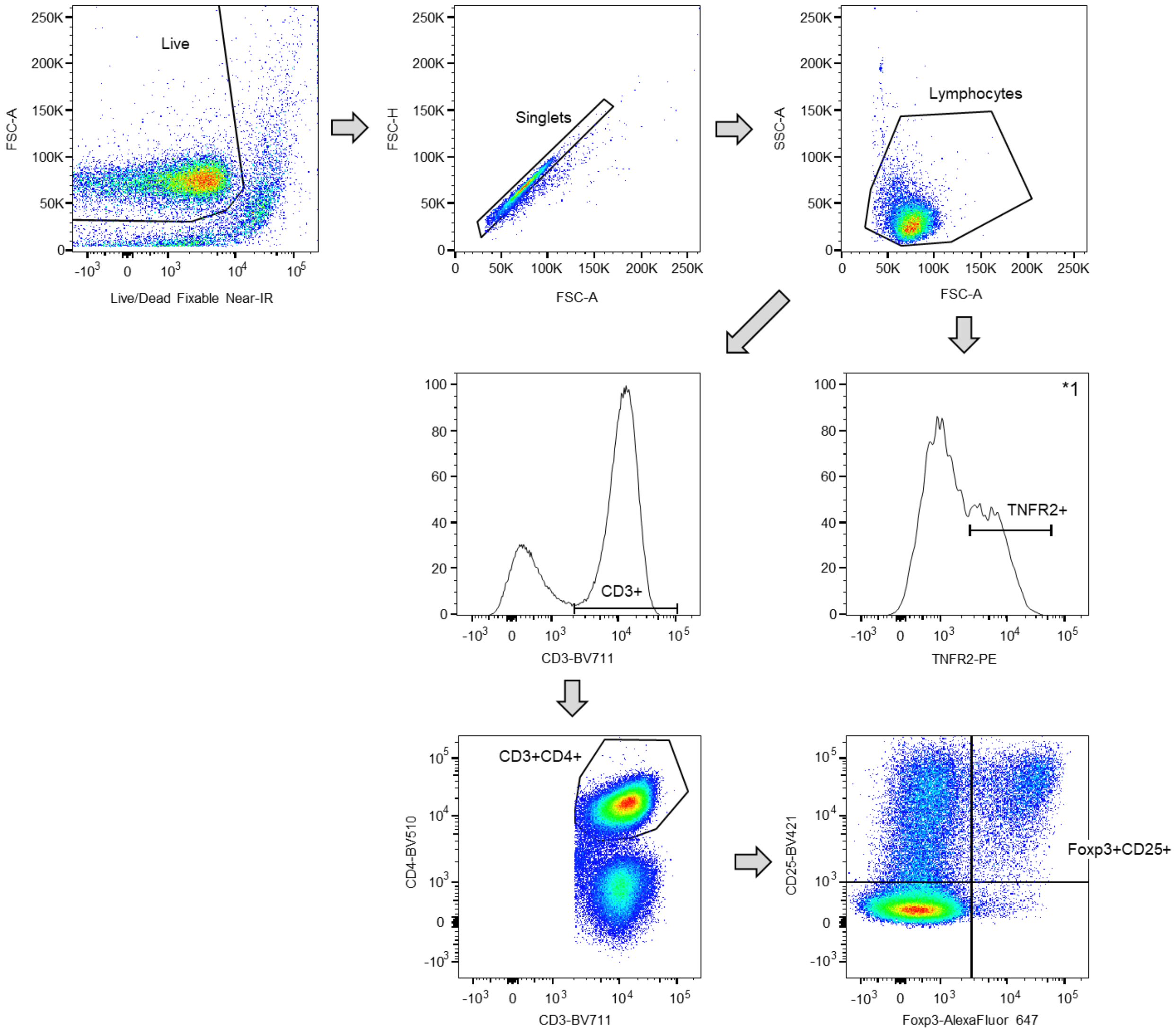


**Fig. S17. Gating strategy for analyzing peripheral blood mononuclear cells.** TNFR2 expression in lymphocytes in the presence of 100 ng/mL TNFα was used for gating for clarity and the same gating was used for all conditions (*1).

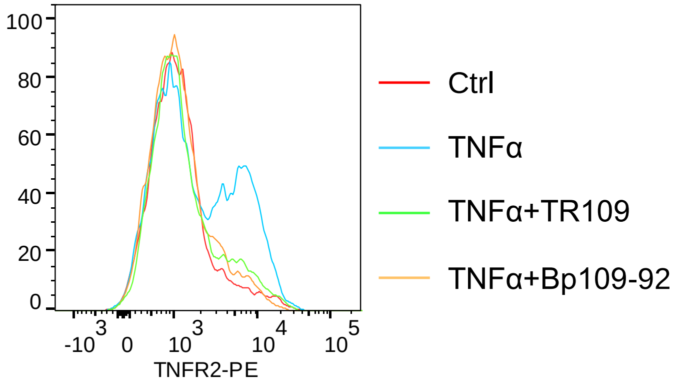


**Fig. S18. CD3⁺ cells analyzed by TNFR2 expression.** Peripheral blood mononuclear cells were cultured for 48 h and the cells were analyzed using flow cytometry. Red, no stimulation (Ctrl); cyan, in the presence of 100 ng/mL TNFα (TNFα); green, in the presence of 500 ng/mL TR109 and TNFα (TNFα+TR109); orange, in the presence of 500 ng/mL Bp109-92 and TNFα (TNFα+Bp109-92).

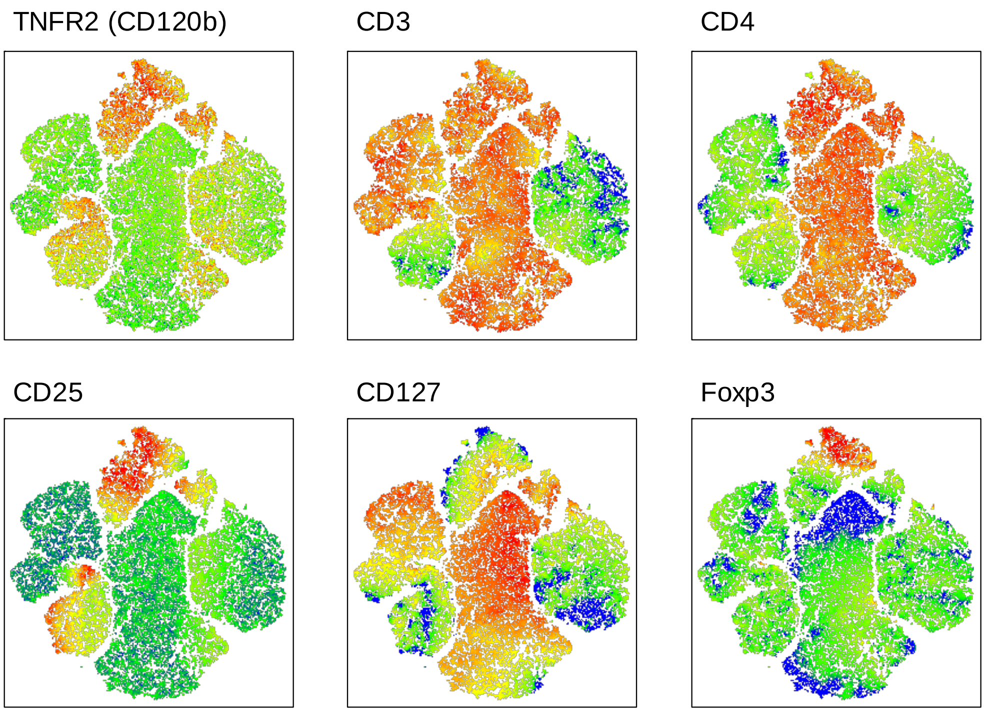


**Fig. S19. TNFR2 expression in relation with several T cell markers found in cluster analysis of PBMC.** The whole PBMC population in eight conditions (10000 counts per condition; see Fig. 4G,H) were clustered by Fast interpolation-based t-SNE algorithm using five parameters (CD3-BV711, CD4-BV510, CD25-BV421, CD127-PerCP-Cy5.5, Foxp3-AlexaFluor 647). Each panel shows the expression of the labeled cellular markers. Color gradient from blue to red indicates low to high expression. For the whole population, TNFR2 expression correlated well with CD25 expression. $CD3^{+}CD4^{+}TNFR2^{+}$ cells were characteristic of low expression of CD127. Most $Foxp3^{+}$ cells were present in the $TNFR2^{+}$ population reflecting correlation of TNFR2 expression with CD25 and CD127.

**Table S1. TNFR2 mutants used for epitope mapping. (separate file)**

**Table S2. Kinetic parameters of TNFR2 binding to antibodies[a].**

| Antibody | $k_{on}$ ($10^6$/Ms) | $k_{off}$ ($10^{-3}$/s) | $K_D$ (nM) |
|---|---|---|---|
| TR45 | 0.22 | 4.1 | 19 |
| TR94[b] | 0.28 | 2.6 | 9.5 |
| TR96 | 0.58 | 0.24 | 0.42 |
| TR92 | 0.87 | 3.5 | 4.0 |
| TR109 | 0.27 | 1.5 | 5.6 |
| Bp109-45 | 0.42 | 0.55 | 1.3 |
| Bp109-92 | 3.1 | 0.48 | 0.16 |
| Bp109-94 | 0.56 | 0.39 | 0.69 |
| Bp109-96 | 0.81 | 0.33 | 0.41 |
| Bp45-92 | 0.70 | 3.1 | 4.5 |
| Bp45-94 | 0.58 | 0.69 | 1.2 |
| Bp45-96 | 1.2 | 0.30 | 0.26 |
| Bp94-92 | 6.2 | 0.61 | 0.098 |
| Bp94-96 | 1.3 | 0.25 | 0.20 |
| Bp96-92 | 1.2 | 0.35 | 0.30 |

[a] Fitting for 1:1 binding kinetics.

[b] Parameters for TR94 are not accurate due to poor fitting.

**Table S3. cryo-EM data collection and image processing of Bp109-92–TNFR2-MBP complex.**

| Dataset | Bp109-92–TNFR2-MBP complex |
|---|---|
| EMDB accession code | EMD-34871 |
| PDB accession code | 8HLB |
| **Data collection and processing** | |
| Magnification | 60,000 |
| Voltage (kV) | 300 |
| Electron exposure ($e^{-}/Å^2$) | 60 |
| Defocus range (μm) | −0.5 to −2.0 |
| Pixel size (Å) | 1.088 |
| Symmetry imposed | C1 |
| Micrographs used (no.) | 4,160 |
| Initial particle images (no.) | 2,218,071 |
| Final particle images (no.) | 100,391 |
| Map resolution (Å) | 3.63 |
| FSC threshold | 0.143 |
| **Refinement** | |
| Initial model used (PDB code) | 3ALQ |
| Model resolution (Å) | 3.5/3.6/4.0 |
| FSC threshold | 0/0.143/0.5 |
| Model vs. Data CC (mask) | 0.75 |
| (volume) | 0.73 |
| Model composition | |
| Non-hydrogen atoms | 7464 |
| Protein residues | 977 |
| Ligands | 0 |
| R.m.s. deviations | |
| Bond lengths (Å) | 0.006 |
| Bond angles (°) | 0.761 |
| Validation | |
| MolProbity score | 2.05 |
| Clashscore | 10.35 |
| Poor rotamers (%) | 0 |
| Ramachandran plot | |
| Favored (%) | 91.21 |
| Allowed (%) | 8.38 |
| Outliers (%) | 0.41 |
